# Microphysiological vascular malformation model reveals a role of dysregulated Rac1 and mTORC1/2 in lesion formation

**DOI:** 10.1101/2022.09.03.506415

**Authors:** Wen Yih Aw, Crescentia Cho, Hao Wang, Anne Hope Cooper, Elizabeth L. Doherty, David Rocco, Stephanie A. Huang, Sarah Kubik, Chloe P. Whitworth, Ryan Armstrong, Anthony J. Hickey, Boyce Griffith, Matthew L. Kutys, Julie Blatt, William J. Polacheck

## Abstract

Somatic activating mutations of *PIK3CA* are associated with the development of vascular malformations (VMs). Here, we describe a microfluidic model of *PIK3CA*-driven VMs consisting of human umbilical vein endothelial cells (HUVECs) expressing *PIK3CA* activating mutations embedded in 3D hydrogels. We observed enlarged and irregular vessel phenotypes, consistent with clinical signatures and concomitant with PI3K-driven upregulation of Rac1/PAK, MEK/ERK, and mTORC1/2 signaling. We observed differential effects between Alpelisib, a PIK3CA inhibitor, and Rapamycin, an mTORC1 inhibitor, in mitigating matrix degradation and vascular network topology. While both drugs are effective in preventing vessel enlargement, Alpelisib suppressed mTORC2-dependent AKT1 phosphorylation and MEK/ERK signaling. Rapamycin failed to reduce MEK/ERK and mTORC2 activity and resulted in vascular hyperbranching, while inhibiting PAK, MEK1/2, and mTORC1/2 signaling mitigates abnormal growth and vascular dilation. Collectively, these findings establish an *in vitro* platform for modeling VMs and confirm a role of dysregulated Rac1/PAK and mTORC1/2 signaling in *PIK3CA*-driven VMs.

## Introduction

Vascular malformations (VMs) are a class of rare genetic disorders associated with localized developmental abnormalities of venous, arterial, capillary, or lymphatic vessels [1, 2]. Histologically, VMs are characterized by a complex of structural lesions with enlarged, irregular lumens that are lined with vascular endothelial cells, surrounded by uneven and disorganized extracellular matrix [1, 3]. These lesions are thought to be congenital, often progress in severity with time, are associated with vascular obstruction and impaired drainage, and can be life-threatening [1–4]. Treatment for VMs, especially for complex lymphatic malformations that occur in multiple body sites such as generalized lymphatic anomaly (GLA) and Gorham-Stout disease (GSD), is difficult and rarely curative. Current interventional approaches rely on surgery, sclerotherapy [5–7], and a limited repertoire of drugs [2, 7, 8].

VMs are caused by somatic mutations in genes involved in vasculature development and the same genes are commonly implicated in malignant tumor angiogenesis [2, 4]. Specific surface receptors including VEGFR3 [9, 10], Notch [11–14], Ephb4 [15, 16], Alk1 [17, 18], and Eng [18, 19] ligands, including BMP9 [20], ANGPT2 [21, 22], and VEGF-C, and signaling proteins, including PIK3CA [23–30], Ras [31, 32], and Raf [33], among others, have been associated with one or more types of malformations. In particular, somatic activating “hotspot” mutations in the *PIK3CA* gene have been identified to be the cause of the majority (∼80%) of cystic lymphatic malformations [25, 26], and several studies have further associated these mutations with GLA, capillary malformations, and venous malformations [23, 24, 27, 28, 30]. *PIK3CA* encodes the catalytic p110α subunit of phosphatidylinositol-3-kinase (PI3K), a component of PI3K lipid kinases that is involved in cell proliferation, migration, contractility, survival, and metabolism regulation [34]. In response to receptor tyrosine kinase signaling, PI3K catalyzes the production of phosphatidylinositol 3,4,5-bisphosphate (PtdIns(3,4,5)P3 or PIP3) in the plasma membrane [34]. PIP3 subsequently recruits Pleckstrin homology (PH) domain-harboring proteins including PDK1, mTORC2, and AKT to the plasma membrane [34–36], triggering phosphorylation of AKT by PDK1 and mTORC2 and activation of AKT1 and mTORC1 signaling networks [34, 36]. In addition to controlling cell growth and proliferation through mTOR signaling, PI3K also regulates cell polarization and migration through interactions with RhoA/Rac1 signaling pathways [37, 38].

Several *in vitro* and *in vivo* animal models have been developed to study the pathobiology and to assess treatment efficacy in *PIK3CA*-related VMs. It has recently been shown that clonal expression of *PIK3CA* activating mutation in the lymphatic endothelium promotes vessel overgrowth and the formation of VMs in mice [27, 28, 30, 39–41], demonstrating a causative role of *PIK3CA* activating mutations in the pathogenesis of VMs. Characterization of these mouse- and patient-derived lymphatic endothelial cells revealed consistent molecular alterations associated with dysregulated PI3K/AKT/mTOR signaling, including inhibition of tumor suppressor tuberin (TSC2), and phosphorylation and hyperactivation of AKT1 and mTORC1/2 signaling. These signaling changes are associated with increased cell size, cell proliferation, and angiogenesis [26, 30, 40]. Despite these established cellular phenotypes, the respective contributions of cellular, molecular, and mechanical factors to disease progression are difficult to isolate and control *in vivo*, and how each of these associated signaling axes might contribute to VMs remains unclear.

Identification of altered PI3K/mTOR signaling has led to the use of mTOR and PI3K inhibitors in treating VMs. Rapamycin (Sirolimus), an mTOR inhibitor, is the most commonly used drug for treating VMs [30, 42]. Recent studies from animal models of *PIK3CA*-mutated breast cancer and *PIK3CA*-driven lymphatic malformations led to successful repurposing of a PIK3CA specific inhibitor (Alpelisib) in treating lymphatic malformations [41, 43]. Treatment with Alpelisib was shown to restore healthy cellular phenotypes and prevent development of and promote regression of vascular lesions in mouse models and in many patients [41, 43]. In contrast, treatment with Rapamycin may be effective at restricting the growth of lesions but appears to have limited efficacy in complete lesion remission [41, 43]. Although both drugs can be effective in treating VMs, the cellular processes that govern growth inhibition and reversal of enlarged vessels remain largely unknown.

Here, we generated human umbilical vein endothelial cells (HUVECs) expressing a PIK3CA activating mutation (*PIK3CA*^*E542K*^) and incorporated PIK3CA^E542K^ HUVECs in 3D vascular networks embedded in physiologic ECM within microfluidic devices. We demonstrate that mutant endothelial cells are hyperproliferative and develop vascular networks with complex, irregular, and enlarged structures, closely resembling clinical phenotypes. These phenotypes were driven by endothelial cell-autonomous hyperactivation of PI3K signaling and were associated with increased cell proliferation and matrix proteolysis in mutant vascular networks. We observed that broad spectrum inhibition of endothelial cell proliferation and matrix proteolysis mitigated lesion formation but did not rescue alterations in vascular morphology. In agreement with clinical data, we demonstrated that Alpelisib is more effective than Rapamycin in preventing the development of malformed vasculature as Rapamycin prevented vascular enlargement but resulted in vascular hyperbranching. We investigated the signaling effects of Alpelisib inhibition and demonstrated that in contrast to Rapamycin, Alpelisib led to suppression of both mTORC2-dependent AKT1 phosphorylation and MEK/ERK signaling in PIK3CA^E542K^ cells. We further identified Rac1/PAK and MEK/ERK activation as key effectors of PIK3CA hyperactivation and utilize our model to demonstrate that targeting PAK, MEK/ERK, and mTORC1/2 signaling are effective in improving malformed vessel structure.

## Results

### Generation and characterization of endothelial cells expressing *PIK3CA* activating mutations

To investigate the effects of *PIK3CA*-activating mutations on vascular endothelial cells, we generated HUVECs expressing *PIK3CA*^*E542K*^*-IRES-GFP* or *PIK3CA*^*E545K*^*-IRES-GFP, PI3K* activating mutations detected in patients with VMs syndromes [23–30], by lentiviral transduction. As controls, we infected HUVECs with the same lentiviral vector encoding GFP or wildtype PIK3CA (PIK3CA^WT^, Fig. S1–S2). GFP expression was used to determine transduction efficiency for each cell line (Fig. S2A). Western blot analysis of phospho-AKT1 confirmed that PI3K/AKT signaling was upregulated in HUVECs expressing *PIK3CA*-activating mutations (Fig. S1C). Immunostaining with VE-cadherin and phalloidin demonstrated distinct cytoskeletal and adhesive phenotypes, with cells expressing PIK3CA^E542K^ or PIK3CA^E545K^ characterized by diminished and discontinuous VE-cadherin adhesions, increased formation of actin stress fibers, and decreased cortical actin localization (Fig. S1A, B). Consistent with previous reports [39], we observed that HUVECs expressing *PIK3CA* mutations were larger in area than controls (Fig. S1A, B). While there are no significant changes in the percent population of senescence cells in PIK3CA^E542K^ or PIK3CA^E545K^ when compared to controls, we observed elevated beta-galactosidase activity in HUVECs that were larger in cell size (Fig. S2B).

### Microfluidic model that recapitulates clinical phenotypes of VMs

While overall these phenotypes are consistent with previous reports on the effects of expressing *PIK3CA* activating mutations on the cell shape and cytoskeletal organization [39], it is unclear how these changes relate to the pathophysiology and clinical presentation of VMs, including disorganized vessel organization and cyst formation. To relate these cellular phenotypic changes to microvascular topology, we generated 3D vascular networks by co-culturing HUVECs and stromal human lung fibroblasts (HLFs) within fibrin hydrogels (Fig. 1A). To visualize vascular network formation for each cell type over time, we imaged the GFP channel using laser scanning confocal microscopy every 24hrs. Strikingly, while GFP and PIK3CA^WT^ cells formed interconnected networks that are well-distributed in 3D, PIK3CA^E542K^ and PIK3CA^E545K^ cells formed irregular vascular networks with expanded vessel diameter that are akin to histological hallmarks of vascular lesions in patients (Fig. 1A-C). The lack of organization in vascular networks was accompanied by an increase in the projected area covered by PIK3CA^E542K^ and PIK3CA^E545K^ cells within tissue constructs when compared to controls (Fig. 1D). Notably, when imaging the fibrin matrix, we observed the presence of cavities that enlarge over time in mutant vascular networks (Fig. 1D, S3). Close inspection of spaces void of ECM revealed that these cavities were composed of dilated vessel enveloped by a continuous layer of endothelial cells (Fig. 1C). The formation of voids coincides with increased levels of secreted proteases in the conditioned media of PIK3CA^E542K^ cells (Fig. S4), implicating a role of dysregulated matrix proteolysis in mutant cells.

**Figure 1.**
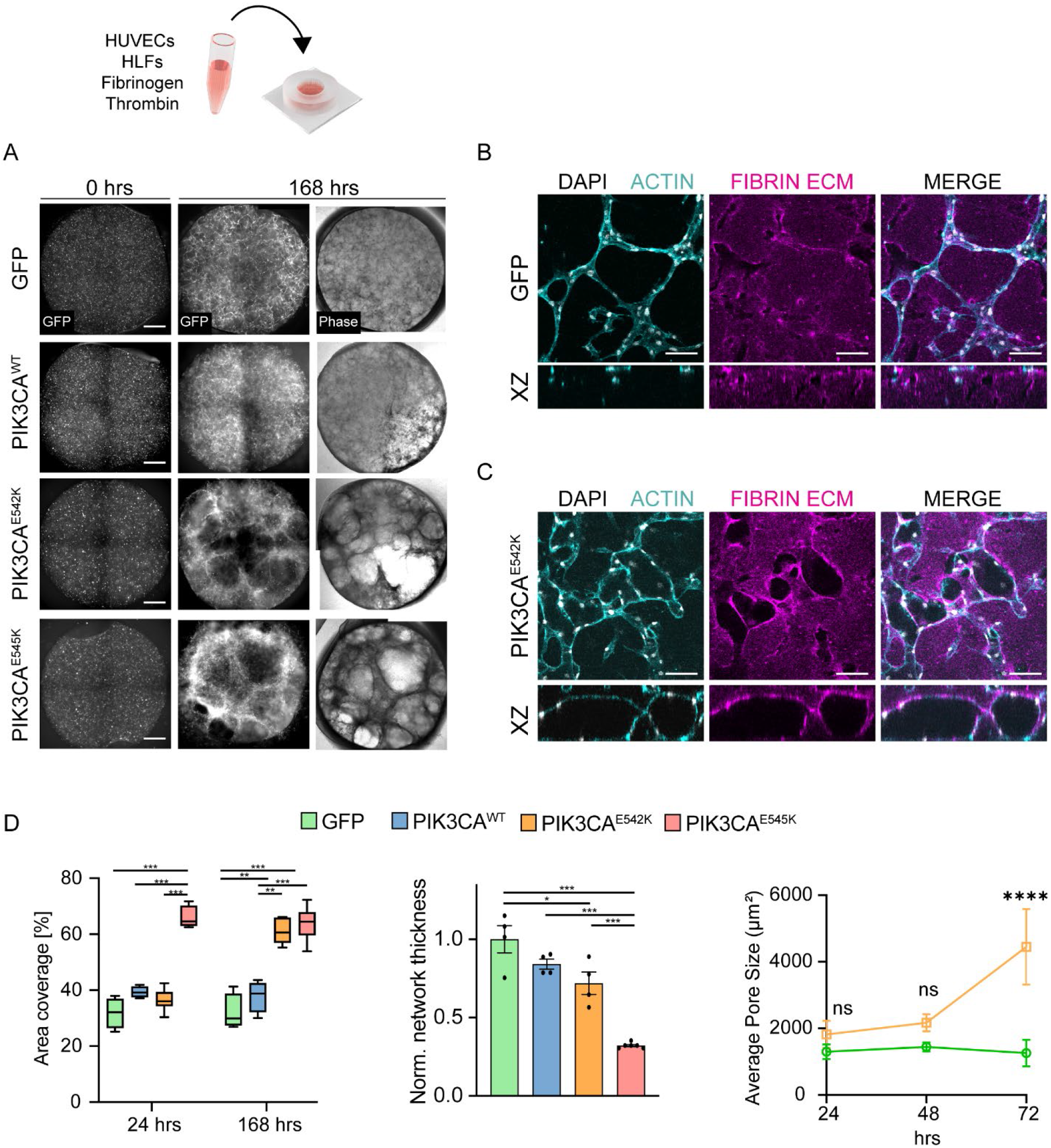
3D culture of endothelial cells expressing constitutively active *PIK3CA* isoforms recapitulates irregular and enlarged vessel phenotypes associated with vascular malformations. (a) Control cells and endothelial cells expressing *PIK3CA* activating mutations cultured in 3D fibrin matrices and imaged at 0 and 168 hours after seeding. Scale bar = 1000 μm. (b, c) Representative confocal images of control and PIK3CA^E542K^ vascular networks. Vascular networks were fixed 3 days after seeding. Scale bar = 100 μm. (d) Quantification of images in (a). (n = 3 to 5 independent experiments; mean ± s.d; One-way ANOVA with Tukey post test, \**p* ≤ 0.05; \*\**p* ≤ 0.01; \*\*\**p* ≤ 0.001; \*\*\*\**p* ≤ 0.0001).

### Endothelial cell-autonomous contributions to pathologic network formation

We next sought to evaluate the role of stromal-endothelial cell interactions in the development of malformed vessel structures observed in 3D culture. Using a multi-channel microfluidic device [44], we seeded HUVECs and stromal fibroblasts in separate spatial compartments and quantified the effects of compartmentalized stromal-vascular network coculture on vessel morphology (Fig. 2A). Consistent with the results when stromal and endothelial cells were embedded within the same hydrogel, we observed abnormal growth and increased vessel dilation in PIK3CA-mutant vascular networks when compared to controls (Fig. 2B-C), suggesting that direct interaction with stromal cells is not essential for initiating vascular malformations. In addition, complete removal of stromal human lung fibroblasts from the culture system modestly reduced PIK3CA^E542K^ vascular volume but did not prevent the development of vascular lesions as quantified by the total volume of fibrin-void cavities (Fig. 2B-C). Together, these data suggest that while malformation growth may be regulated by non-autonomous signaling between endothelial cells and surrounding cell types, the development of VMs are driven mostly by cell-autonomous PI3K activation in endothelial cells.

**Figure 2.**
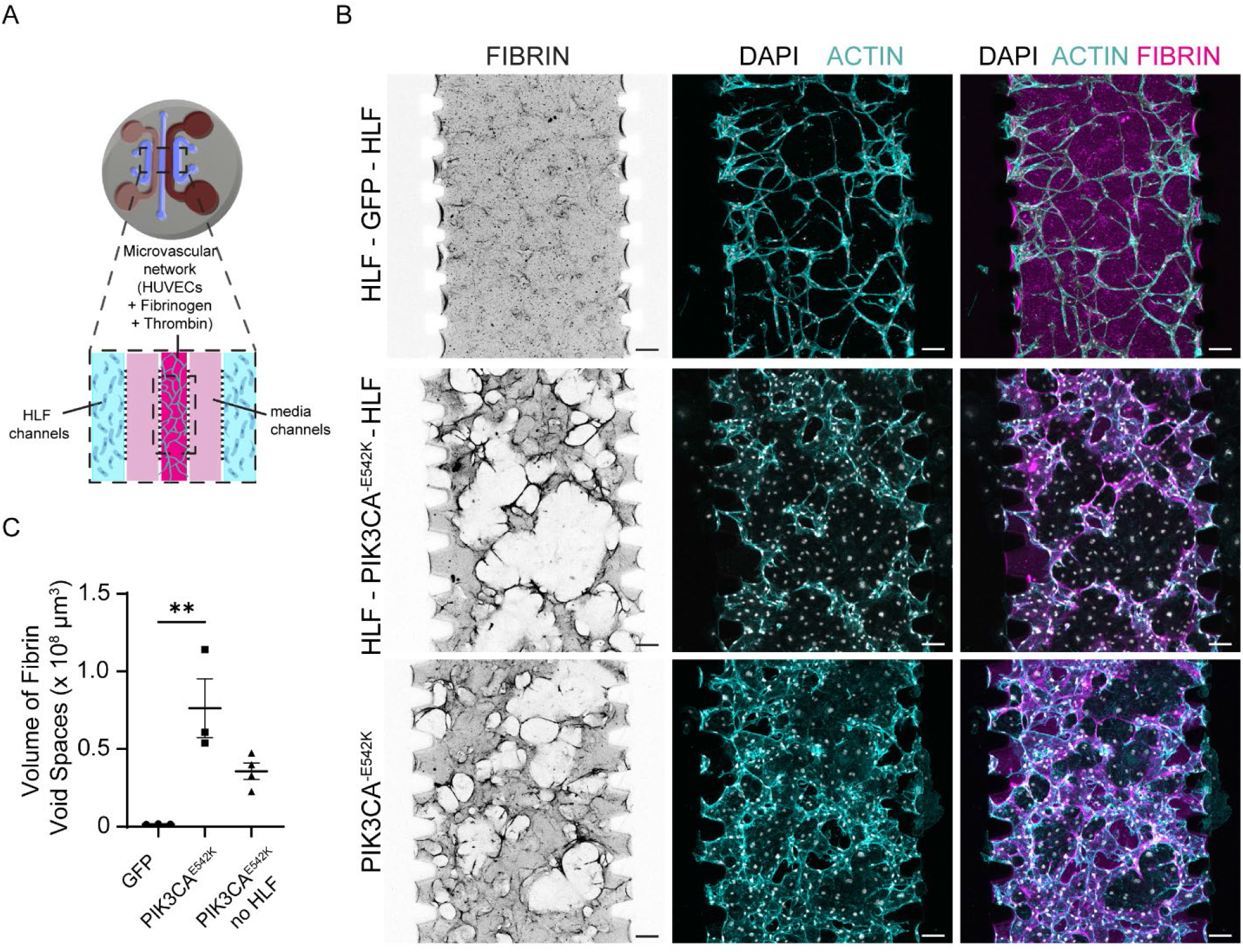
PIK3CA^E542K^ functions largely cell-autonomously in driving vascular overgrowth and vessel enlargement. (a) Schematic of microfluidic device used for culturing endothelial cells and human lung fibroblasts (HLFs) in separate spatial compartments. (b) Confocal images of GFP and PIK3CA^E542K^ vascular networks labeled with Fibrin (magenta), DAPI (white), and Actin (cyan). Vascular networks were generated by co-culturing endothelial cells and HLFs in different spatial compartments (HLF-endothelial cells-HLF) or by monoculture of PIK3CA^E542K^ cells in all three compartments. Vascular networks were fixed 3 days after seeding. Scale bar = 100 μm. (c) Volumetric quantification of void spaces in fibrin hydrogel (n = 3; mean ± s.e.m; One-way ANOVA with Tukey post test, \**p* ≤ 0.05; \*\*\**p* ≤ 0.001; \*\*\*\**p* ≤ 0.0001).

To gain further insights into the mechanisms driving vessel enlargement, we performed live cell imaging of vascular networks cultured in fluorescently-labeled fibrin hydrogel. This allowed us to capture dynamics of cell-cell and cell-ECM interactions in control and PIK3CA^E542K^ cells over 86hr period of vascular network formation (Fig. 3, Supplementary Movie 1). Consistent with results from fixed samples, GFP cells formed an interconnected vascular network with a relatively uniform distribution of vessel diameter (Fig. 3A, Supplementary Movie 1). We observed local ECM remodeling as reflected by increased fibrin intensity in the vicinity of control vascular network (Supplementary Movie 1). In comparison to control, we found excessive ECM remodeling and degradation which are associated with lumen expansion in PIK3CA^E542K^ vascular network (Fig. 3B, Supplementary Movie 1). We conclude from these data that PIK3CA-driven VMs results from abnormal proliferation and dysregulated matrix degradation of mutant endothelial cells.

**Figure 3.**
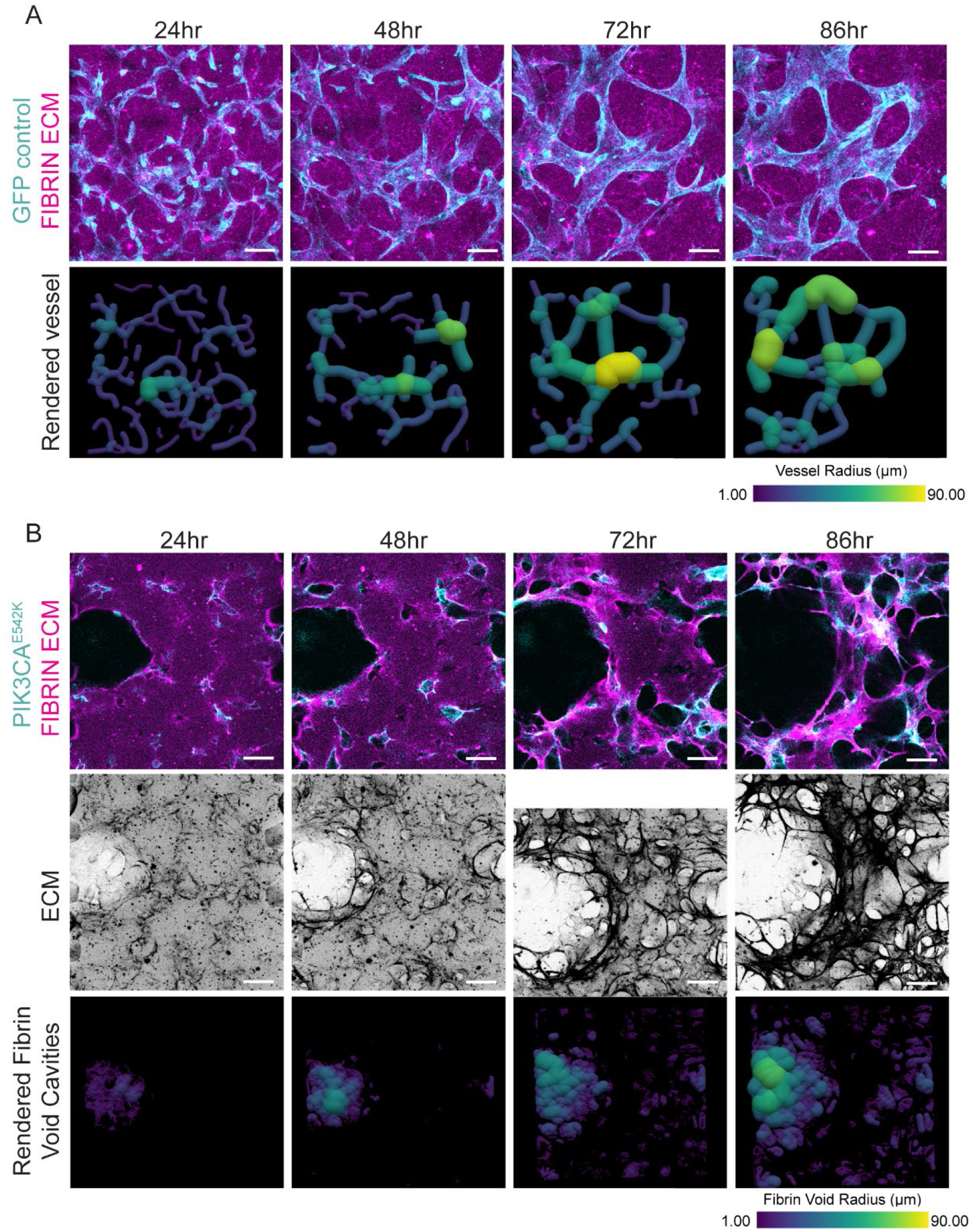
Temporal progression of vascular malformation-like phenotypes in PIK3CA^E542K^ vascular networks. (a) Time-lapse confocal images (maximum intensity projections) of GFP control endothelial cells (cyan) cultured in fibrin ECM (magenta) and corresponding volume rendering of segmented vascular networks. Scale bar = 100 μm. (b) Single plane time-lapse images of PIK3CA^E542K^ cells (cyan) cultured in fibrin ECM (magenta) (top), maximum intensity projections of fibrin ECM (mid), and corresponding 3D rendering of segmented ECM void spaces. Scale bar = 100 μm.

### Inhibition of cell proliferation and matrix degradation attenuates VM phenotypes

To study the role of endothelial cell proliferation in vascular enlargement, we inhibited cell proliferation by treating PIK3CA^E542K^ cells with mitomycin C. After 3 days of culture, the number of cells within mutant vascular networks was similar to that observed in the control networks (Fig. 4A, B).

**Figure 4.**
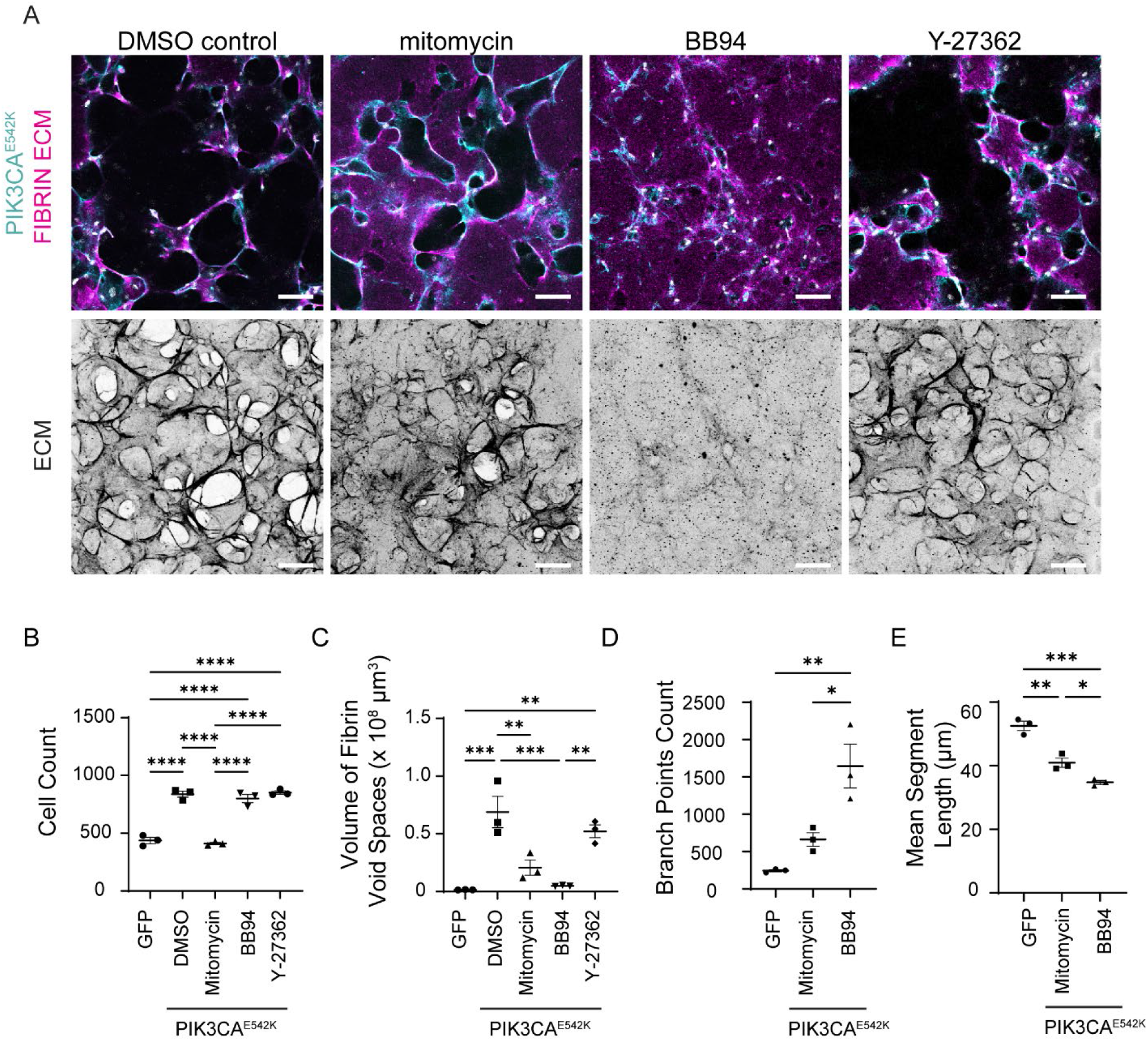
Inhibition of endothelial cell proliferation and matrix proteolysis mitigates but is not sufficient to rescue malformed vascular architecture. (a) Representative images of PIK3CA^E542K^ vascular networks cultured for 3 days with DMSO, BB94 (1μM), or Y-27362 (10μM). For mitotic inhibition, PIK3CA^E542K^ cells were pretreated with 0.01 mg/mL mitomycin-C prior to culturing in fibrin hydrogel. Scale bar = 100 μm. (b-e) Cell count (b), volumetric quantification of ECM (c), vessel branching (d) and segment length (e) of GFP control and PIK3CA^E542K^ vascular networks (n = 3; mean ± s.e.m; One-way ANOVA with Tukey post test, \**p* ≤ 0.05; \*\**p* ≤ 0.01; \*\*\**p* ≤ 0.001; \*\*\*\**p* ≤ 0.0001).

Visual examination of ECM and quantification of vascular structure showed a reduction in ECM remodeling and volume of vascular lesions with mitotic inhibition (Fig. 4A, C). Despite the attenuated severity in ECM degradation, treatment with the mitotic inhibitor did not fully rescue the vascular topology driven by PIK3CA^E542K^ expression (Fig. 4A, D, E). As compared to GFP control, mitotically inhibited PIK3CA^E542K^ vascular networks exhibited enlarged vessels and increased vascular branching (Fig. 4A, D). A similar hyperbranching morphology was observed upon treatment with batimastat (BB-94), a broad-spectrum metalloproteinase inhibitor (Fig. 4A-E), implying that inhibition of cell proliferation or matrix degradation individually is not sufficient to rescue healthy microvascular topology. As elevated Rho kinase signaling is implicated in the development of cerebral cavernous malformations (CCM) [45–47], a type of VMs in the brain and spinal cord, we tested whether the development of VMs in our model required Rho kinase activity by inhibiting PIK3CA^E542K^ cells with Y27362, a Rho-kinase inhibitor. While we observed modest attenuation in ECM remodeling with Y27362 treatment, the morphologies of mutant vascular networks treated with Y27362 were indistinguishable from those of the untreated control (Fig. 4A-C), suggesting that targeting RhoA activity and cellular contractility is not effective in preventing *PIK3CA*-driven VMs. Collectively, these results highlight that proliferation, matrix degradation, cytoskeletal dynamics, and tissue contractility are dysregulated in PIK3CA^E542K^ vascular networks, but targeting them individually does not rescue the phenotype, suggesting that intervention is necessary upstream of these cellular phenotypes.

### Inhibition of PIK3CA rescues network topology, while inhibition of mTOR signaling results in network hyperbranching

Rapamycin and Alpelisib have both been demonstrated to be effective in treating *PIK3CA*-driven VMs patients [30, 42, 43]. Because a comparative study in a *PIK3CA* mouse model demonstrated that Alpelisib is more effective than Rapamycin in improving organ abnormalities [41], we sought to test and compare their efficacies in treating VMs in our tissue engineered human vascular network model (Fig. 5, S5A, C). We observed reduced cell proliferation and drastic reduction in ECM proteolysis as measured by the volume of fibrin void cavities with either Rapamycin or Alpelisib treatments (Fig. 5A, C, D). While both drugs prevented the development of vascular lesions as measured by the volume of void spaces in fibrin hydrogel (Fig. 5D), we observed significant differences in treatment responses as manifested by vascular branching patterns (Fig. 5A, E, F). In particular, PIK3CA^E542K^ networks treated with Rapamycin exhibited a vascular hyperbranching phenotype, as demonstrated by the formation of network with shorter vessel segments (Fig. 5A, E) and the increased number of vessel branches and end points (Fig. 5A, F). Consistent with a previous report [41], treatment with Alpelisib effectively reduced mTORC2 activity as evidenced by reduced AKT1 phosphorylation at Ser473 by western blot analysis, but Rapamycin was not effective in inhibiting mTORC2 activity (Fig. 5G-I). Further, we observed reduced ERK1/2 phosphorylation in PIK3CA^E542K^ cells treated with Alpelisib, but not with Rapamycin (Fig. 5G-I), suggesting that in addition to mTORC2 signaling, MEK/ERK activity is differentially regulated by these inhibitors. Overall, these results suggest that crosstalk between PI3K and MEK/ERK signaling may contribute to VMs.

**Figure 5.**
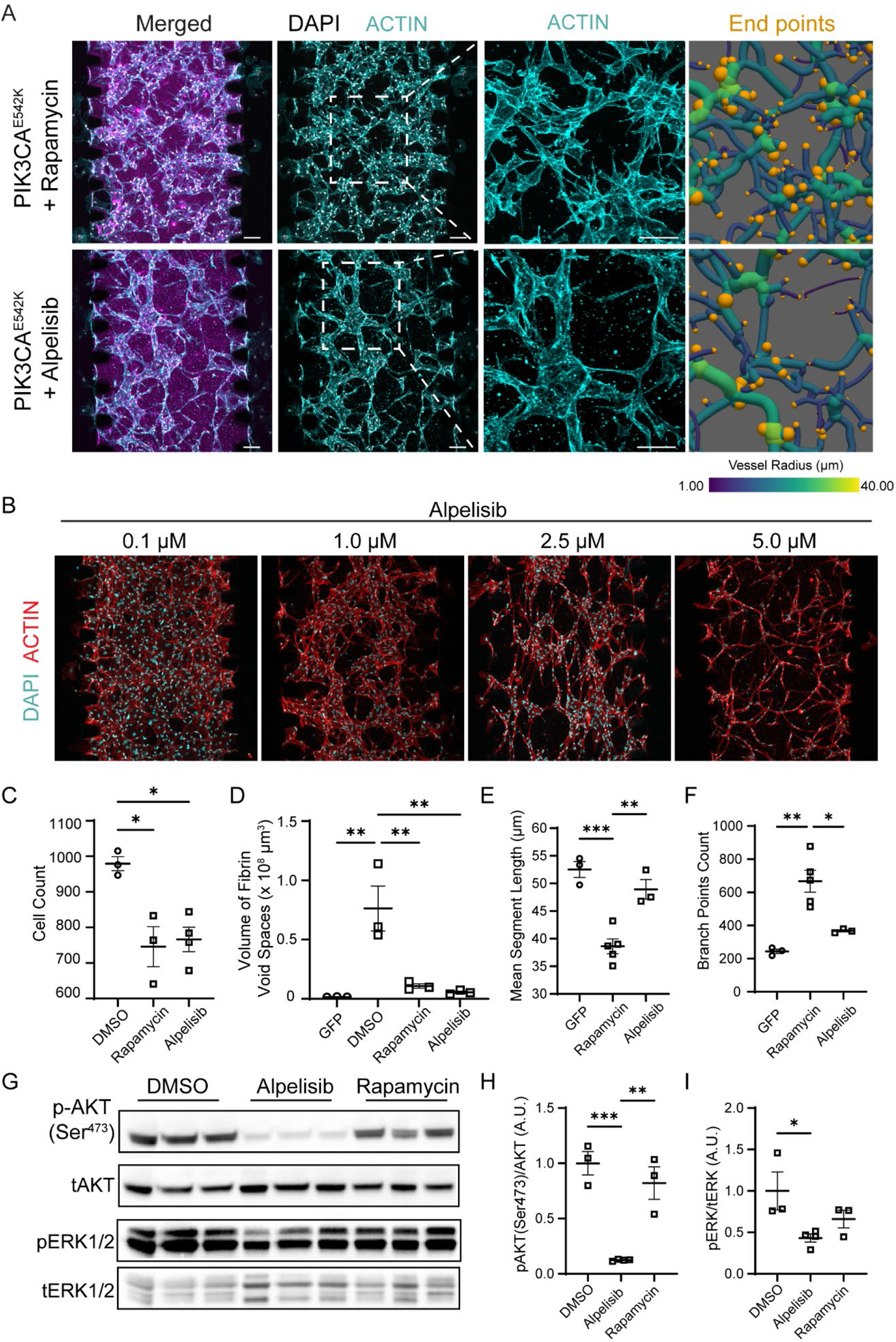
Rapamycin treatment improves vascular malformation but leads to vascular hyperbranching. (a) Representative images of PIK3CA^E542K^ vascular networks treated with Rapamycin (1μM) or Alpelisib (2.5μM) for 3 days. Vascular networks were labeled with fibrin (magenta), DAPI (white), and Actin (cyan). Endpoints (orange) were highlighted in 3D renderings of segmented vessels. Scale bar = 100 μm. (b) Dose-dependent changes in PIK3CA^E542K^ vascular morphology with Alpelisib treatment. (c-f) Quantification of cell number (c), ECM morphology (d), mean vessel length (e), and vessel branching (f) in GFP control, Rapamycin treated PIK3CA^E542K^ vascular networks, or Alpelisib treated PIK3CA^E542K^ vascular networks (n ≥ 3; mean ± s.e.m; One-way ANOVA with Tukey post test). (g-i) Western blot and quantification of relative p-AKT1(Ser473) and p-ERK1/2(Thr202/Tyr204) levels in PIK3CA^E542K^ cells treated with DMSO, Rapamycin, or Alpelisib (n ≥ 3; mean ± s.e.m; One-way ANOVA with Tukey post test, \**p* ≤ 0.05; \*\**p* ≤ 0.01; \*\*\**p* ≤ 0.001).

### Targeting elevated MEK1/2 signaling with Trametinib reduces vascular lesions but results in vascular hyperbranching

To test the relevance of MEK/ERK activation in PIK3CA-driven vascular malformations, we treated PIK3CA^E542K^ vascular network with Trametinib (Fig. 6, S5C), a MEK1/2 kinase inhibitor that has proven to be effective in improving VMs in patients with activating *MAP2K1* or *KRAS* mutations and in *KRAS*-driven mouse models of GSD [48–50]. Inhibition with Trametinib reduced protease secretion by PIK3CA^E542K^ cells (Fig S4). Western blot analysis revealed that Trametinib blocked ERK1/2 phosphorylation but not mTORC2 dependent AKT1-Ser473 phosphorylation (Fig. 6A, C-D). While reduced ERK1/2 phosphorylation is distinctive of Trametinib treatment (Fig. 6A, D), Trametinib-treated PIK3CA^E542K^ vascular networks displayed a reduced vascular lesions and hyperbranched vascular network morphology that is comparable to the inhibition of mTORC1 with rapamycin (Fig. 6B, E-F). These results indicate that MEK/ERK and mTORC1 signaling pathways share overlapping roles in the regulation of vascular growth as inhibition of either MEK/ERK or mTORC1 signaling alleviates vascular overgrowth but results in abnormal hyperbranched vascular networks.

**Figure 6.**
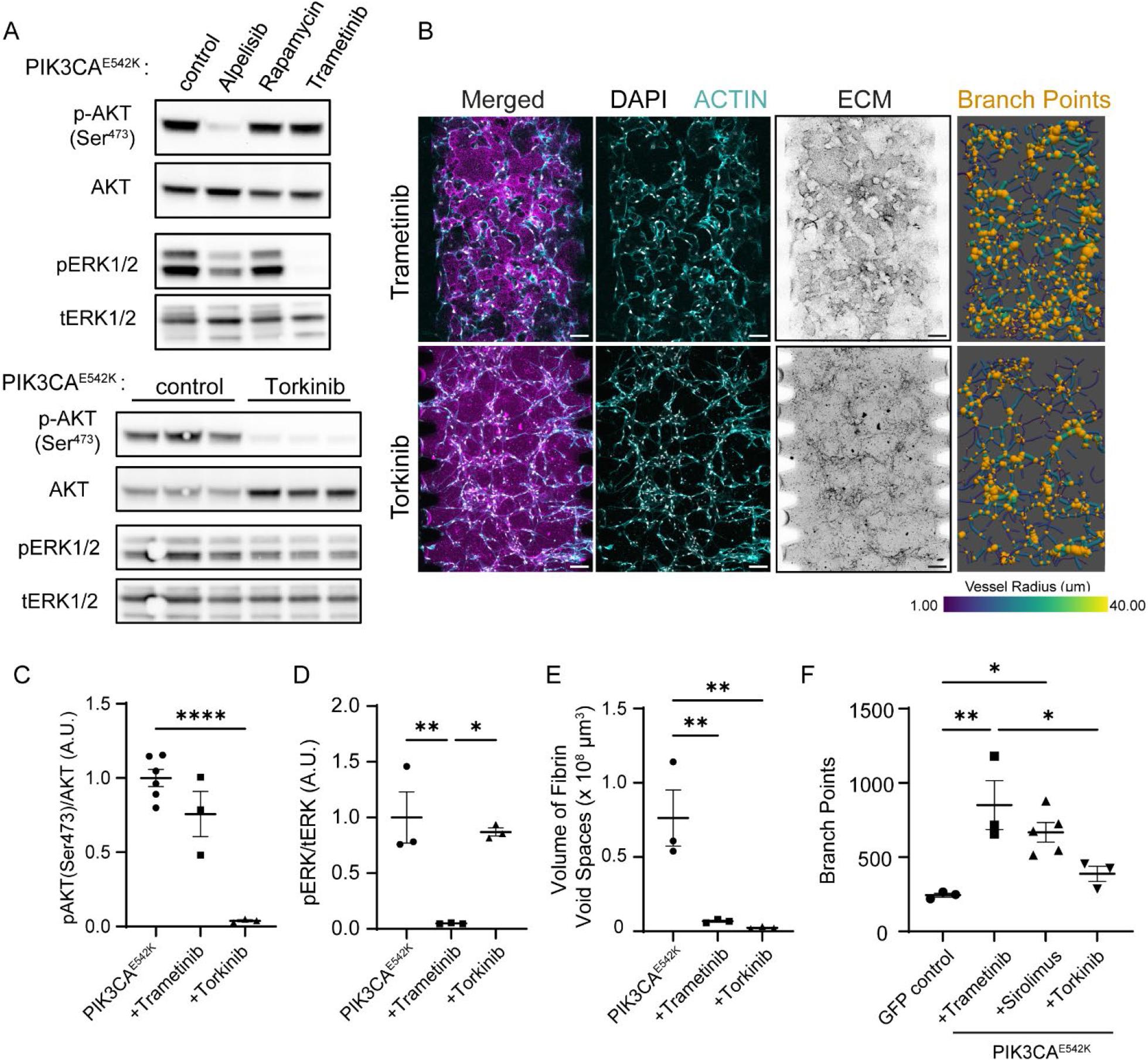
Treatment with Trametinib (MEK1/2 inhibitor) and Torkinib (mTORC1/2 inhibitor) improves and rescues vascular phenotypes in PIK3CA^E542K^ vascular networks, respectively. (a) Western blot analysis of relative p-AKT1(Ser473) and p-ERK1/2(Thr202/Tyr204) in PIK3CA^E542K^ cells treated with the indicated inhibitors. (b) Representative confocal images of PIK3CA^E542K^ treated with Trametinib (MEK1/2 inhibitor, 1μM) or Torkinib (mTORC1/2 inhibitor, 5μM) for 3 days. Scale bar = 100 μm. (c-d) Quantification of relative p-AKT1 and p-ERK1/2 level in (a) (n ≥ 3; mean ± s.e.m; One-way ANOVA with Tukey post test). (e-f) Quantification of matrix (e) and vessel branching morphology (f) in (b) (n ≥ 3; mean ± s.e.m; One-way ANOVA with Tukey post test, \**p* ≤ 0.05; \*\**p* ≤ 0.01; \*\*\*\**p* ≤ 0.0001).

### Dual inhibition of mTORC1 and mTORC2 rescues network topology

Given the known role of mTORC2 signaling in regulating actin cytoskeletal dynamics [51] and our observation that rapamycin treatment led to incomplete suppression of mTORC2 signaling, we postulated that vascular hyperbranching is a consequence of dysregulated mTORC2 signaling. To test this, we treated PIK3CA^E542K^ vascular networks with Torkinib, an ATP-competitive mTORC1/2 inhibitor [52, 53], and compared the resulting vascular morphology to that of the control GFP or Rapamycin treated PIK3CA^E542K^ vascular network (Fig. 6F). In contrast to Rapamycin, treatment with Torkinib reduced phosphorylation of AKT1-Ser473 (Fig. 6A, C) and prevented the development of vascular lesions and hyperbranching phenotypes (Fig. 6B, E-F). Further, when compared to Trametinib, treatment with Torkinib had a greater effect on suppressing protease secretion (Fig. S4), implicating a role of mTORC1/2 signaling in matrix degradation. Collectively, these results are consistent with previous observations demonstrating that mTORC2 regulates endothelial cell sprouting and migration partially through regulation of actin cytoskeleton and focal adhesion dynamics [54], and indicates a combined role of mTORC1-dependent overgrowth and mTORC2-dependent cytoskeletal defects in the development of malformed vascular networks.

### Elevated Rac1 activity contributes to VM

Previous studies have demonstrated the role of dysregulated small GTPase signaling and MEK/ERK activation in the development of complex lymphatic malformations [32, 50, 55–58]. One potential explanation for increased MEK/ERK signaling could be elevated small GTPase activity in PIK3CA^E542K^ cells. To explore this hypothesis, we examined the effect of Alpelisib treatment on Ras, Rac, and Cdc42-GTPase activities in PIK3CA^E542K^ cells and found that treatment with Alpelisib resulted in reduced Rac-GTP level (Fig. S5B). Consistent with this observation, we found higher levels of ERK1/2 phosphorylation and Rac-GTP in PIK3CA^E542K^ cells when compared to control (Fig. 7A-E). As Rac1-GTPase plays a critical role in mediating actin polymerization and membrane protrusion at the leading edge of migrating cells, we performed time-lapse imaging and investigated the effect of *PIK3CA* mutation on endothelial cell migration (Fig. 7F-J, Supplementary Movie 2).

**Figure 7.**
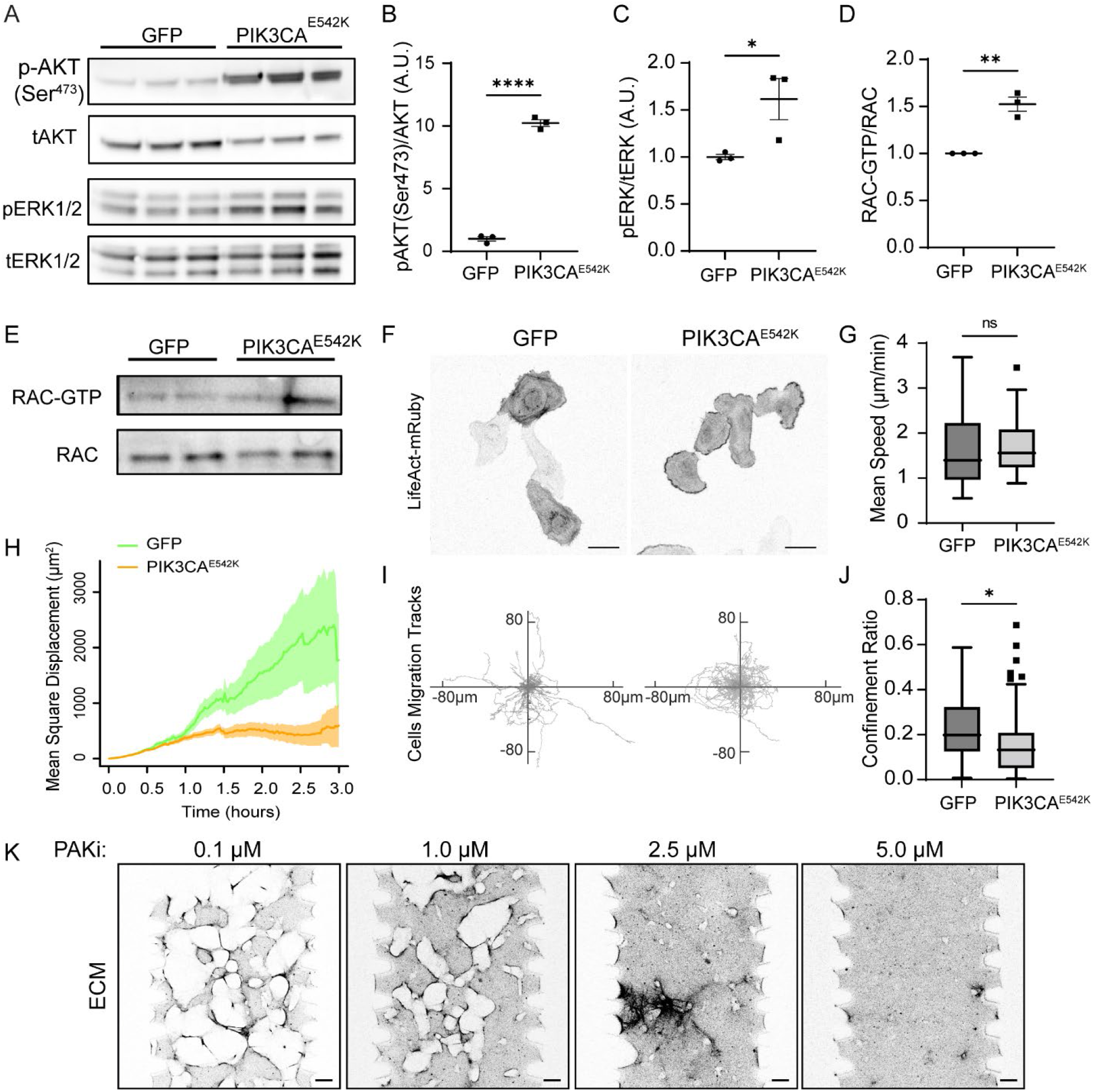
Endothelial cells expressing *PIK3CA* mutation display molecular signatures of excessive Rac1 and MEK/ERK signaling. (a-c) Western blot analysis (a) and quantification of relative p-AKT1(Ser473) (b) and p-ERK1/2(Thr202/Tyr204) (c) levels in serum-starved GFP control and PIK3CA^E542K^ endothelial cells (n = 3; mean ± s.e.m; two-tailed unpaired Student t-test, \**p* ≤ 0.05; \*\*\*\**p* ≤ 0.0001). (d-e) Quantification (d) and western blot (e) of active Rac1 level in serum-starved GFP control and PIK3CA^E542K^ cells. Active Rac1 was isolated using a GST-tagged p21-binding domain of PAK1 (n = 3; mean ± s.e.m; two-tailed unpaired Student t-test, \*\**p* ≤ 0.01). (f) Representative images of GFP and PIK3CA^E542K^ cells expressing LifeAct-mRuby, cultured in 2D. Note the presence of actin-rich membrane ruffles at the periphery of PIK3CA^E542K^ cells. Scale bar = 50 μm. (g-j) Mean migration speed (g), mean square displacement (h), migration trajectories (i) and confinement ratios as defined by net-displacement over total travelled distance (j) of GFP and PIK3CA^E542K^ cells (n = 57 and 60 tracks, Tukey box and whisker, two-tailed unpaired Student t-test, \**p* ≤ 0.05). (k) Representative confocal images of ECM surrounding PIK3CA^E542K^ vascular networks treated with 0.1μM to 5.0μM of PAK inhibitor, PF-3758309. Scale bar = 100 μm.

Consistent with cellular phenotypes associated with Rac1 activation [59, 60], we found that PIK3CA^E542K^ cells produced persistent actin-rich membrane ruffles around cell edges (Fig. 7F, Supplementary Movie 2). Further, we observed clear differences in cell migration behavior, where control cells migrate more persistently than cells expressing *PIK3CA*^*E542K*^ mutation (Fig. 7I). To quantify this effect, we calculated the mean migration speed (Fig. 7G), mean square displacement (Fig. 7H), and the confinement ratio of control and mutant cells (Fig. 7J). While we observed no significant difference in migration speed, PIK3CA^E542K^ cells showed significantly reduced mean square displacement and confinement ratios as defined by net-displacement over total travelled distance (Fig. 7H-J), demonstrating that persistent cell migration was inhibited with constitutive PI3K activation in endothelial cells expressing *PIK3CA* mutation. We further tested the effect of PAK1 inhibition on PIK3CA^E542K^ vascular networks. Treatment with PAK inhibitor PF-3758039 resulted in a dose-dependent reduction of vascular lesions as measured by volume of fibrin void spaces in PIK3CA^E542K^ vascular networks (Fig. 7K, S5C, S6B, C). Notably, we observed changes in 3D vascular network morphology where vascular networks appeared sheet-like and flattened with higher doses of PAK inhibitions (Fig. S6A). Interestingly, all doses of PAK inhibition led to a similar reduction in endothelial cell number, indicating that these dose-dependent morphology changes are not solely mediated by alteration in endothelial cell proliferation (Fig. S6C). Together, these results confirm that both MEK/ERK and Rac-GTPase activities are upregulated with PI3K activation, and suggest that excessive Rac/PAK and MEK/ERK signaling may be targetable in *PIK3CA*-driven VMs.

## Discussion

Here, we generated vascular networks with HUVECs expressing a *PIK3CA* activating mutation (Fig. S1–S2) and demonstrated that the mutant endothelial cells develop into dilated vascular networks with complex and irregular lumens that resemble clinical hallmarks observed in patients with vascular malformations (Fig. 1–3). These dramatic phenotypes observed in 3D culture are striking in comparison to the relatively mild differences observed for the same cells in 2D cell culture (Fig. S1–S2). We found that vessel enlargement and cyst development are driven largely by endothelial cell-autonomous defects that are associated with increased endothelial cell proliferation and dysregulated cell-mediated matrix proteolysis and tissue mechanics (Fig. 2, 4, S4, S5). Using our model, we tested the efficacies of Rapamycin, a mTORC1 inhibitor, and Alpelisib, a PIK3CA specific inhibitor, in treating VMs (Fig. 5, S5C). We analyzed the resulting vascular network geometry in 3D and discovered that while both inhibitors prevented vessel enlargement and improved vascular network morphology, treatment with Rapamycin resulted in vessel hyperbranching (Fig. 5).

We observed that Rapamycin was ineffective in suppressing phosphorylation of the mTORC2 target, AKT1 Ser473 (Fig. 5G-H), when compared to Alpelisib. mTOR serine/threonine kinase is a downstream effector of PI3K signaling and functions through two multiprotein complexes, mTORC1 and mTORC2 [61]. While mTOR is known to regulate endothelial cell proliferation and angiogenesis [62–64], the specific manner in which mTORC1 and mTORC2 contribute to the cellular processes involved in angiogenesis remains to be defined. Using our model, we showed that inhibition of mTORC1 with Rapamycin resulted in vascular hyperbranching (Fig. 5A, F).

Notably, dual inhibition of mTORC1/2 rescued vessel hyperbranching (Fig 6B, F), indicating a specific role of mTORC2 in regulating vessel length and branchpoint selection during vessel formation.

*In vitro* studies have demonstrated a role of mTORC2 in controlling actin cytoskeleton rearrangement and cell spreading in fibroblasts [51]. More recently, studies of mTOR signaling in neurons further showed that mTORC2 plays a vital role in controlling neuron size, morphology, and synaptic function [65, 66]. This is thought to be mediated through regulation of actin cytoskeleton rearrangements and protrusion dynamics via mTORC2-Tiam1-Rac1 signaling [65]. Notably, our data show that both mTORC2 and Rac1 activities are elevated in endothelial cells expressing *PIK3CA*^*E542K*^ mutation (Fig. 7A-E). At the cellular level, we observed increased cell spreading, excess membrane protrusion, and cell migration defects that are consistent with Rac1 activation (Fig. 7F-J). Collectively, our results point to excess mTORC2 and Rac1/Pak signaling leading to dysregulated actin cytoskeletal dynamics and endothelial cell motility in cells expressing *PIK3CA* activating mutations.

The majority of mutations identified in VMs occur within key components of two major growth factor-induced signaling pathways, PI3K-AKT-mTOR and RAS-MAPK-ERK [2, 4]. While both pathways regulate vascular development and tumor angiogenesis, a mechanistic understanding of how activating mutations in PI3K and MAPK signaling lead to the formation of cysts associated with vascular malformations remains elusive. Our study found increased levels of phosphorylated-ERK1/2 in PIK3CA^E542K^ cells when compared to controls (Fig. 7A). Further, treatment of mutant cells with Alpelisib, a PIK3CA specific inhibitor, decreased phospho-ERK1/2 levels (Fig. 5G, 6A), suggesting a role of dysregulated MAPK signaling in *PIK3CA*-driven VMs. Treatment with Trametinib, a MEK1/2 inhibitor, prevented vessel enlargement but resulted in hyperbranched vessel morphology that is similar to vascular networks treated with rapamycin (Fig. 6B, F). This is unsurprising, as MAPK scaffolds are known to regulate several nodes of mTORC1 signaling [67], and the level of mTORC2 dependent AKT1-Ser473 phosphorylation remains unchanged with MEK1/2 inhibition (Fig. 6A, C). Collectively, our findings demonstrate a functional redundancy in the ERK1/2 and mTORC1 signaling pathways in regulating endothelial cell proliferation and vessel enlargement, and that inhibition of cell growth through ERK1/2 or mTORC1 alone is not sufficient to rescue cytoskeletal perturbations that are regulated by mTORC2/Rac1/PAK signaling. These findings are in line with the two-hit model of cerebral cavernous malformations (CCM), where loss of function mutations of the CCM complex that result in increased MEKK3-ERK5-KLF signaling and a subsequent gain of function mutation in *PIK3CA* signaling synergize to promote aggressive CCM lesions [68, 69].

The tissue engineered vascular malformation model described here enables *in vitro* vascular disease modeling, comparative drug studies, and drug-specific pathway discovery. We demonstrate the model’s utility in elucidating cross-talk between PI3K/AKT/mTOR, MEK/ERK, and Rac1/PAK signaling, and illustrate changes in underlying vessel morphology in response to targeted inhibition of these signaling axes. The development of VMs is a complex and multifactorial processes, involving cytoskeletal reorganization, ECM remodeling, proliferation, and potentially changes in endothelial cell contractility. Understanding how each of the PI3K-associated signaling axes contribute to these distinct cell behaviors identifies key signaling nodes relevant not only to *PIK3CA*-driven VMs, but also other types of VMs. We anticipate that the approach described here will motivate the adaptation of vasculature-on-chip models for use in future studies of vascular disease as well as drug screening.

## Acknowledgements

This work was supported by the National Institutes of Health (R35GM142944), the American Heart Association (CDA857738), and by research grants from the University of Pennsylvania Orphan Disease Center in partnership with the Lymphangiomatosis & Gorham’s Disease Alliance and the Lymphatic Malformation Institute, the UNC Computational Medicine Institute, and the North Carolina Biotechnology Center (FLG-3814). W.Y.A. is supported by grants from the CLOVES Syndrome Community and the Eshelman Institute for Innovation. E.L.D., S.A.H., and C.W. acknowledge financial support through the Integrative Vascular Biology Training Program (T32HL69768), and E.L.D. acknowledges support from a Ruth L. Kirchstein predoctoral individual fellowship (F31HL162462). Device fabrication was performed in the Chapel Hill Analytical and Nanofabrication Laboratory, CHANL, a member of the North Carolina Research Triangle Nanotechnology Network, RTNN, which is supported by the National Science Foundation (ECCS-2025064), as part of the National Nanotechnology Coordinated Infrastructure, NNCI.

## Author Contributions

W.Y.A, M.L.K., J.B., and W.J.P. designed the experiments. W.Y.A., C.C, D.R., and E.L.D. created the lentiviral constructs and PIK3CA mutant HUVECs. C.C. conducted the mixed HUVECs-stromal cells culture experiment. W.Y.A., H.W., A.H.C., E.L.D., C.P.W., and R.A. contributed to vascular network morphometric analysis. H.W. and S.A.H. contributed to microfluidic fabrication and cell migration analysis. S.K. and H.W. contributed to data visualization. A.J.H, B.G, M.L.K., and J.B provided essential advice to the project. W.Y.A and W.J.P. wrote the manuscript with inputs from all authors.

---

**Supplemental Figure 1.**
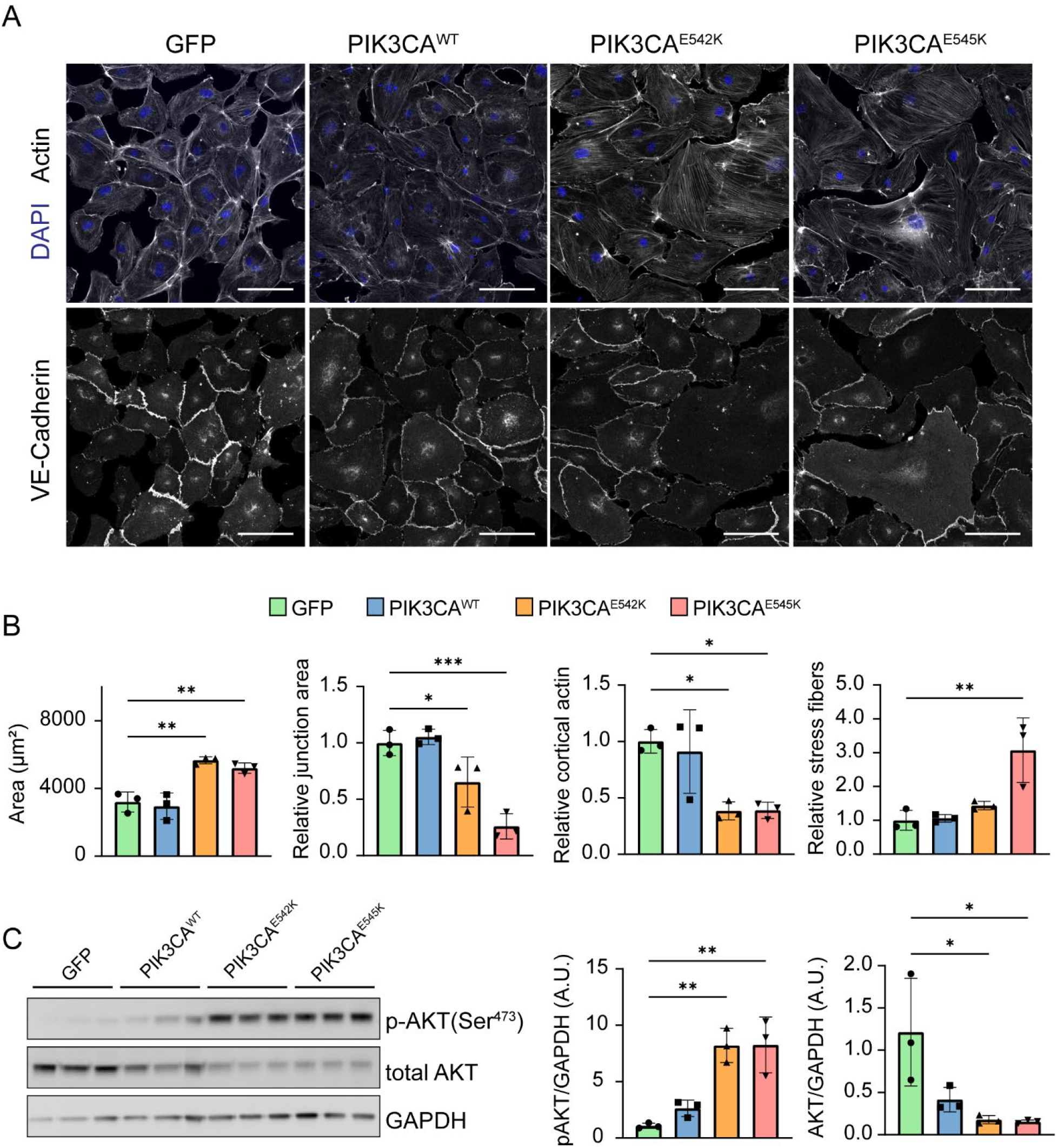
Increased PI3K/AKT activation and altered cytoskeletal organization in endothelial cells expressing *PIK3CA* activating mutations. (a) Representative confocal images of GFP, PIK3CA^WT^, PIK3CA^E542K^, and PIK3CA^E545K^ cells stained for DAPI, Actin, and VE-Cadherin. Scale bar = 100 μm. (b) Quantification of cell area, relative junctional area measured from VE-cadherin staining, relative cortical actin, and stress fiber distributions of GFP, PIK3CA^WT^, PIK3CA^E542K^, and PIK3CA^E545K^ cells. (n = 3; mean ± s.d; One-way ANOVA with Tukey post test, \**p* ≤ 0.05; \*\**p* ≤ 0.01; \*\*\**p* ≤ 0.001). (c) Western blot and quantification of p-AKT1(Ser473) and total-AKT1 levels in GFP, PIK3CA^WT^, PIK3CA^E542K^, and PIK3CA^E545K^ cells (n = 3; mean ± s.d; One-way ANOVA with Tukey post test).

**Supplemental Figure 2.**
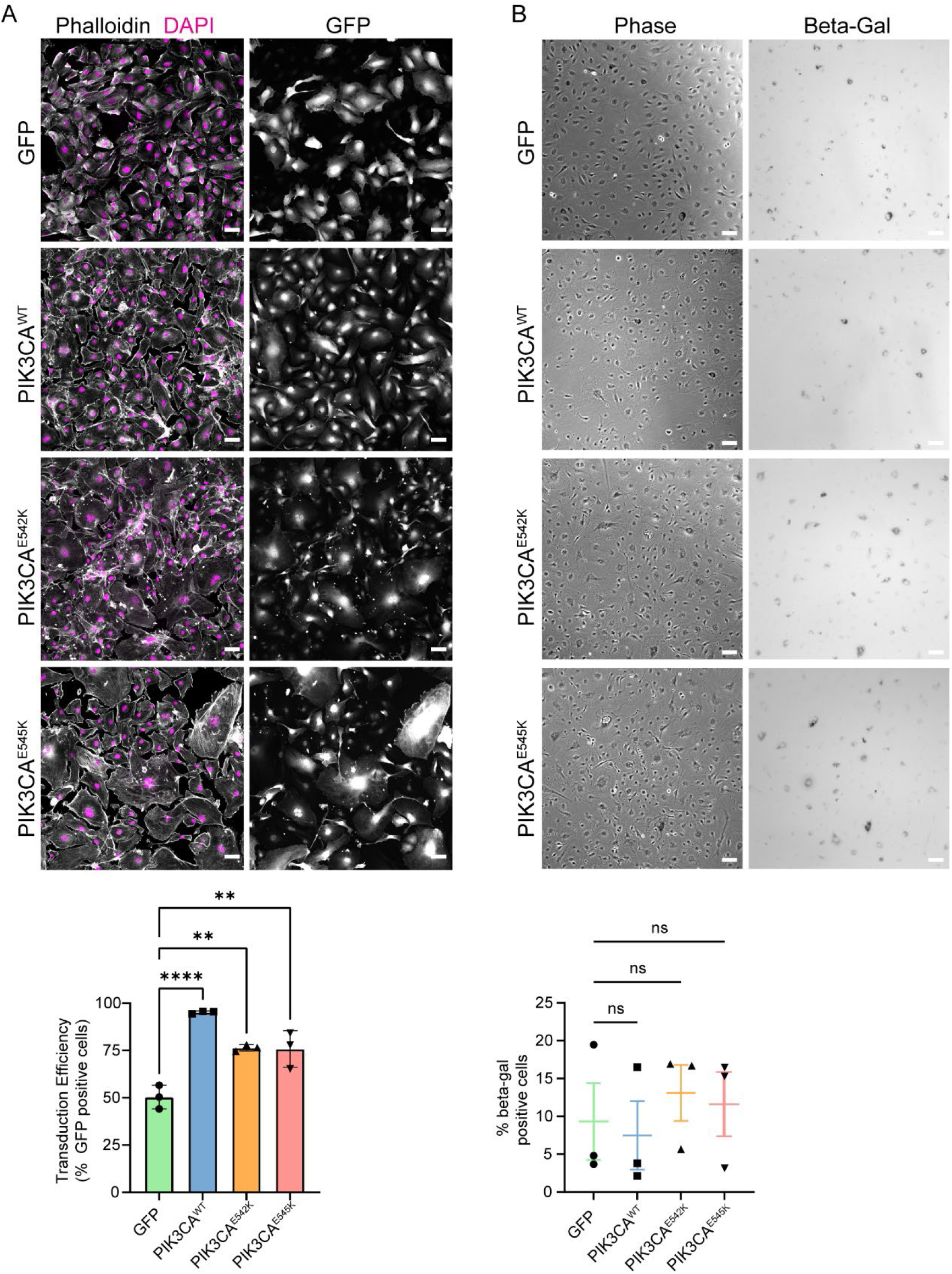
Characterization of transduction efficiency and senescence-associated beta-galactosidase markers in control and mutant endothelial cells. (a) Confocal images of GFP, PIK3CA^WT^, PIK3CA^E542K^, and PIK3CA^E545K^ cells stained for DAPI, Actin, and GFP. Scale bar = 100 μm. Quantification of transduction efficiency of control and cells expressing PIK3CA activating mutations (n = 3; mean ± s.d; One-way ANOVA with Tukey post test, \*\**p* ≤ 0.01; \*\*\*\**p* ≤ 0.0001). (b) B-galactosidase staining and quantification for senescent cells. Scale bar = 100 μm (n = 3; mean ± s.e.m; One-way ANOVA with Tukey post test, ns = not significant).

**Supplemental Figure 3.**
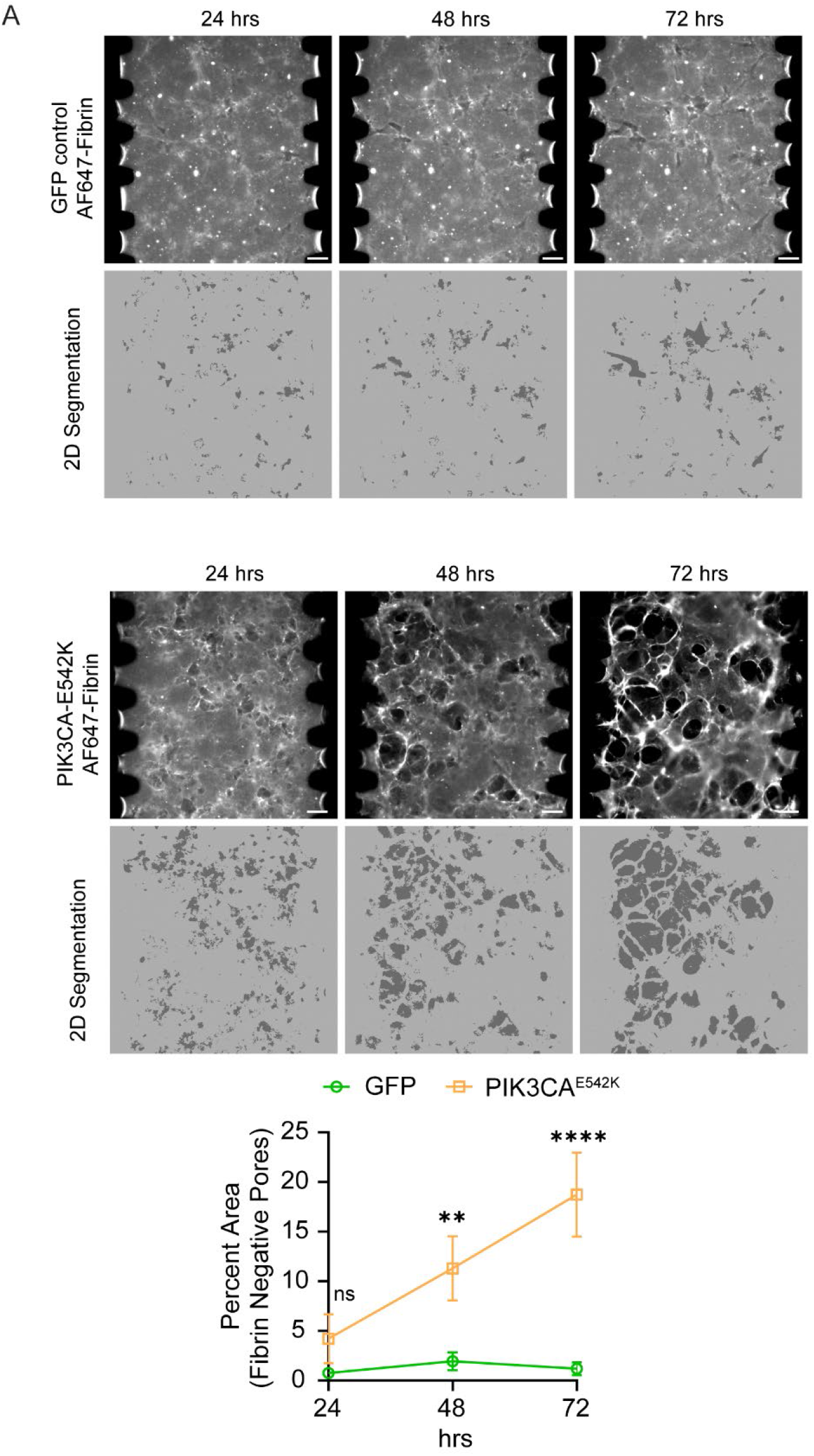
Dysregulated matrix remodeling and increased fibrinolysis in PIK3CA^E542K^ vascular networks. (a) Wide-field images of control and PIK3CA^E542K^ ECM. Fibrin ECM was fluorescently labeled with AF647-fibrinogen and binary masks for spaces void of fibrin signal were segmented for pore size and percent fibrin negative area quantification. Scale bar = 100 μm (n = 3 for GFP control and 6 PIK3CA^E542K^ vascular network; mean ± s.d; One-way ANOVA with Tukey post test, ns = not significant; \*\**p* ≤ 0.01; \*\*\*\**p* ≤ 0.0001).

**Supplemental Figure 4.**
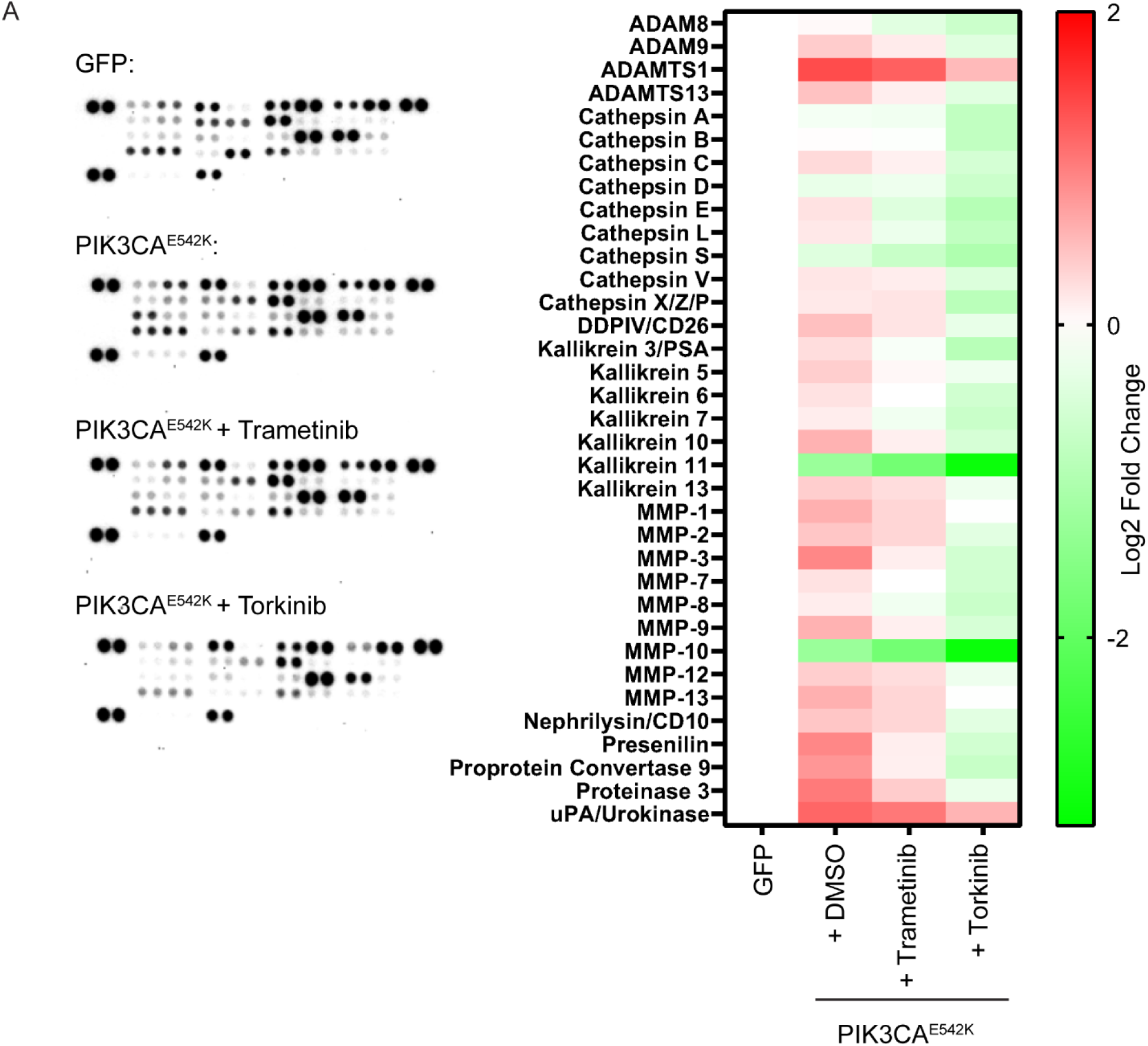
Expression profile of proteases secreted by GFP control treated with DMSO, PIK3CA^E542K^ treated with DMSO, and PIK3CA^E542K^ cells treated with either Trametinib or Torkinib. Endothelial cells were cultured overnight in media with DMSO, 1.0μm Trametinib, or 5.0μm Torkinib. (n = 1)

**Supplemental Figure 5.**
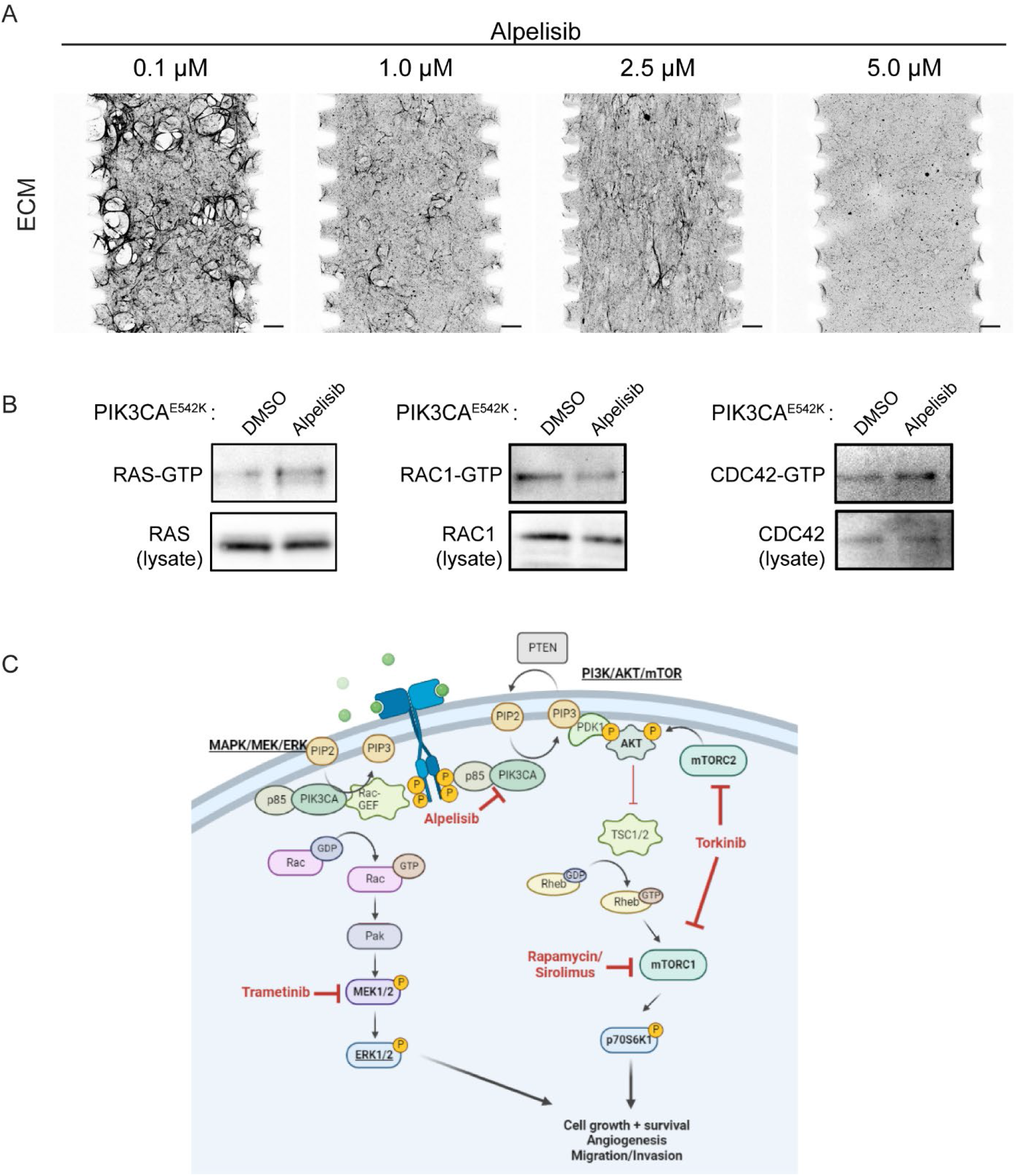
Effects of Alpelisib treatment on PIK3CA^E542K^ vascular networks. (a) Representative ECM images of PIK3CA^E542K^ vascular networks treated with low to high doses of Alpelisib. Scale bar = 100 μm. (b) Active RAS, RAC1, and CDC42 were isolated from PIK3CA^E542K^ cell lysates using GST-Raf-RBD and GST-PBD pull-down. Cells were cultured in a reduced serum condition (0.5% FBS) and were stimulated with complete EGM2 medium with DMSO or 1.0μm Alpelisib for 10 minutes before lysis (n = 1). (c) Schematic representation of dysregulated PI3K/AKT/mTOR and MEK/ERK signaling in PIK3CA mutant cells. Schematic was created with BioRender.com. Tested inhibitors in this study are highlighted in red.

**Supplemental Figure 6.**
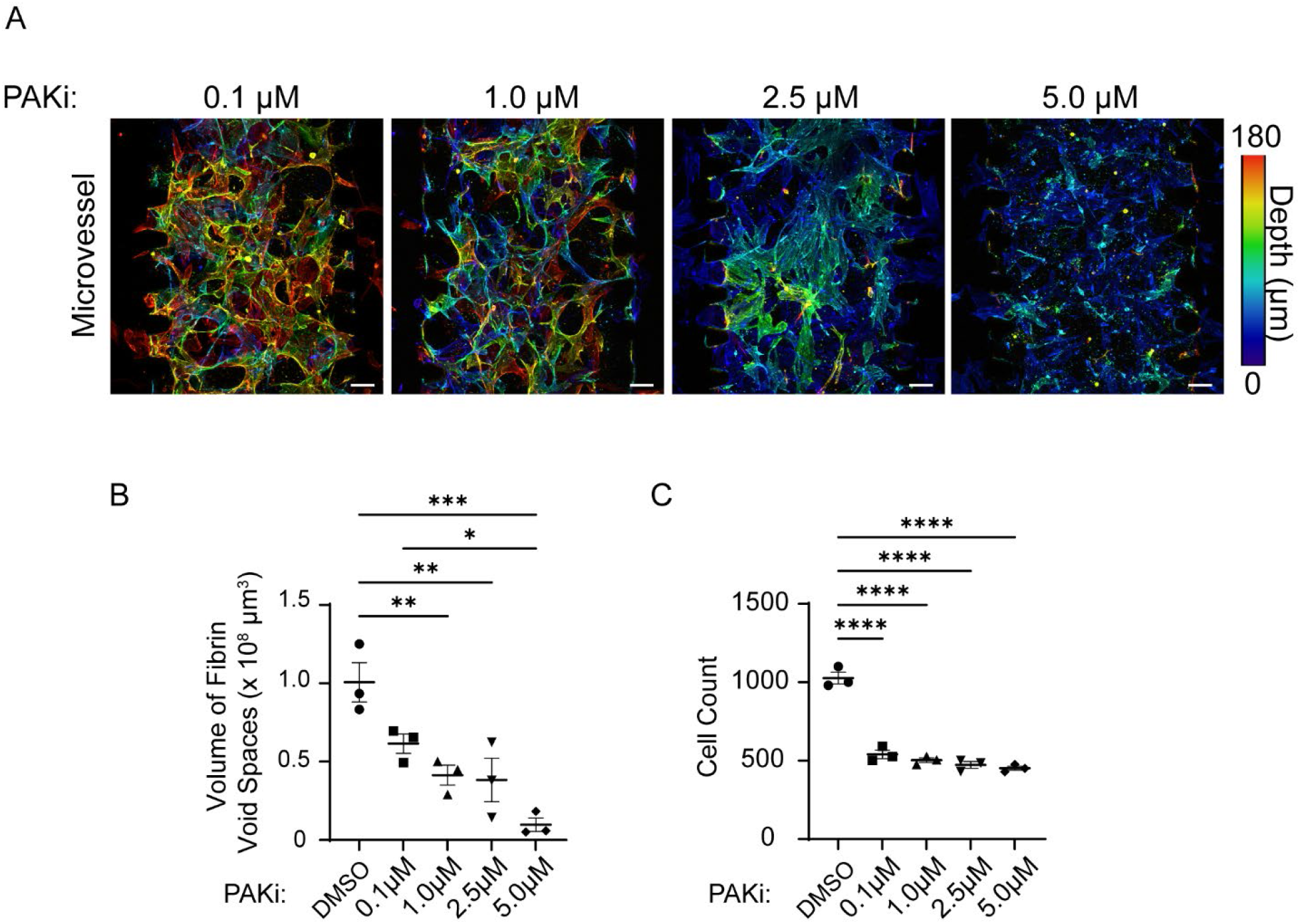
Effects of Pak inhibition on PIK3CA^E542K^ vascular networks. (a) Depth color coded projections of PIK3CA^E542K^ vascular networks treated with 0.1μM to 5.0μM concentration of PAK inhibitor, PF-3758309. Scale bar = 100 μm. (b-c) Quantification of matrix morphology (b) and cell number (c) in PIK3CA^E542K^ vascular networks treated with PAK inhibitor (n = 3; mean ± s.e.m; One-way ANOVA with Tukey post test, \**p* ≤ 0.05; \*\**p* ≤ 0.01; \*\*\**p* ≤ 0.001; \*\*\*\**p* ≤ 0.0001).

**Supplemental Movie 1**. Formation of GFP control and PIK3CA^E542K^ vascular networks over time.

**Supplemental Movie 2**. Example of GFP control and PIK3CA^E542K^ cell membrane ruffling and cell migration.

## References

1. Sadick, M., et al., Vascular Anomalies (Part I): Classification and Diagnostics of Vascular Anomalies. Rofo, 2018. 190(9): p. 825–835.

2. Queisser, A., et al., Genetic Basis and Therapies for Vascular Anomalies. Circ Res, 2021. 129(1): p. 155–173.

3. Cox, J.A., E. Bartlett, and E.I. Lee, Vascular malformations: a review. Semin Plast Surg, 2014. 28(2): p. 58–63.

4. Makinen, T., et al., Lymphatic Malformations: Genetics, Mechanisms and Therapeutic Strategies. Circ Res, 2021. 129(1): p. 136–154.

5. McCafferty, I., Management of Low-Flow Vascular Malformations: Clinical Presentation, Classification, Patient Selection, Imaging and Treatment. Cardiovasc Intervent Radiol, 2015. 38(5): p. 1082–104.

6. Horbach, S.E., et al., Sclerotherapy for low-flow vascular malformations of the head and neck: A systematic review of sclerosing agents. J Plast Reconstr Aesthet Surg, 2016. 69(3): p. 295–304.

7. Kalwani, N.M. and S.G. Rockson, Management of lymphatic vascular malformations: A systematic review of the literature. J Vasc Surg Venous Lymphat Disord, 2021. 9(4): p. 1077–1082.

8. Ng, A.T., R.L. Tower, and B.A. Drolet, Targeted treatment of vascular anomalies. Int J Womens Dermatol, 2021. 7(5Part A): p. 636–639.

9. Karkkainen, M.J., et al., Missense mutations interfere with VEGFR-3 signalling in primary lymphoedema. Nat Genet, 2000. 25(2): p. 153–9.

10. Mendola, A., et al., Mutations in the VEGFR3 signaling pathway explain 36% of familial lymphedema. Mol Syndromol, 2013. 4(6): p. 257–66.

11. Murphy, P.A., et al., Endothelial Notch signaling is upregulated in human brain arteriovenous malformations and a mouse model of the disease. Lab Invest, 2009. 89(9): p. 971–82.

12. ZhuGe, Q., et al., Notch-1 signalling is activated in brain arteriovenous malformations in humans. Brain, 2009. 132(Pt 12): p. 3231–41.

13. Krebs, L.T., et al., Notch1 activation in mice causes arteriovenous malformations phenocopied by ephrinB2 and EphB4 mutants. Genesis, 2010. 48(3): p. 146–50.

14. Davis, R.B., et al., Notch signaling pathway is a potential therapeutic target for extracranial vascular malformations. Sci Rep, 2018. 8(1): p. 17987.

15. Martin-Almedina, S., et al., EPHB4 kinase-inactivating mutations cause autosomal dominant lymphatic-related hydrops fetalis. J Clin Invest, 2016. 126(8): p. 3080–8.

16. Amyere, M., et al., Germline Loss-of-Function Mutations in EPHB4 Cause a Second Form of Capillary Malformation-Arteriovenous Malformation (CM-AVM2) Deregulating RAS-MAPK Signaling. Circulation, 2017. 136(11): p. 1037–1048.

17. Johnson, D.W., et al., Mutations in the activin receptor-like kinase 1 gene in hereditary haemorrhagic telangiectasia type 2. Nat Genet, 1996. 13(2): p. 189–95.

18. Govani, F.S. and C.L. Shovlin, Hereditary haemorrhagic telangiectasia: a clinical and scientific review. Eur J Hum Genet, 2009. 17(7): p. 860–71.

19. McAllister, K.A., et al., Endoglin, a TGF-beta binding protein of endothelial cells, is the gene for hereditary haemorrhagic telangiectasia type 1. Nat Genet, 1994. 8(4): p. 345–51.

20. Wooderchak-Donahue, W.L., et al., BMP9 mutations cause a vascular-anomaly syndrome with phenotypic overlap with hereditary hemorrhagic telangiectasia. Am J Hum Genet, 2013. 93(3): p. 530–7.

21. Wouters, V., et al., Hereditary cutaneomucosal venous malformations are caused by TIE2 mutations with widely variable hyper-phosphorylating effects. Eur J Hum Genet, 2010. 18(4): p. 414–20.

22. Soblet, J., et al., Variable Somatic TIE2 Mutations in Half of Sporadic Venous Malformations. Mol Syndromol, 2013. 4(4): p. 179–83.

23. Kurek, K.C., et al., Somatic mosaic activating mutations in PIK3CA cause CLOVES syndrome. Am J Hum Genet, 2012. 90(6): p. 1108–15.

24. Limaye, N., et al., Somatic Activating PIK3CA Mutations Cause Venous Malformation. Am J Hum Genet, 2015. 97(6): p. 914–21.

25. Luks, V.L., et al., Lymphatic and other vascular malformative/overgrowth disorders are caused by somatic mutations in PIK3CA. J Pediatr, 2015. 166(4): p. 1048-54 e1-5.

26. Osborn, A.J., et al., Activating PIK3CA alleles and lymphangiogenic phenotype of lymphatic endothelial cells isolated from lymphatic malformations. Hum Mol Genet, 2015. 24(4): p. 926–38.

27. Castel, P., et al., Somatic PIK3CA mutations as a driver of sporadic venous malformations. Sci Transl Med, 2016. 8(332): p. 332ra42.

28. Castillo, S.D., et al., Somatic activating mutations in Pik3ca cause sporadic venous malformations in mice and humans. Sci Transl Med, 2016. 8(332): p. 332ra43.

29. Blesinger, H., et al., PIK3CA mutations are specifically localized to lymphatic endothelial cells of lymphatic malformations. PLoS One, 2018. 13(7): p. e0200343.

30. Rodriguez-Laguna, L., et al., Somatic activating mutations in PIK3CA cause generalized lymphatic anomaly. J Exp Med, 2019. 216(2): p. 407–418.

31. Manevitz-Mendelson, E., et al., Somatic NRAS mutation in patient with generalized lymphatic anomaly. Angiogenesis, 2018. 21(2): p. 287–298.

32. Barclay, S.F., et al., A somatic activating NRAS variant associated with kaposiform lymphangiomatosis. Genet Med, 2019. 21(7): p. 1517–1524.

33. Li, D., et al., ARAF recurrent mutation causes central conducting lymphatic anomaly treatable with a MEK inhibitor. Nat Med, 2019. 25(7): p. 1116–1122.

34. Fruman, D.A., et al., The PI3K Pathway in Human Disease. Cell, 2017. 170(4): p. 605–635.

35. Franke, T.F., et al., Direct regulation of the Akt proto-oncogene product by phosphatidylinositol-3,4-bisphosphate. Science, 1997. 275(5300): p. 665–8.

36. Manning, B.D. and A. Toker, AKT/PKB Signaling: Navigating the Network. Cell, 2017. 169(3): p. 381–405.

37. Yang, H.W., et al., Cooperative activation of PI3K by Ras and Rho family small GTPases. Mol Cell, 2012. 47(2): p. 281–90.

38. Campa, C.C., et al., Crossroads of PI3K and Rac pathways. Small GTPases, 2015. 6(2): p. 71–80.

39. di Blasio, L., et al., PI3K/mTOR inhibition promotes the regression of experimental vascular malformations driven by PIK3CA-activating mutations. Cell Death Dis, 2018. 9(2): p. 45.

40. Martinez-Corral, I., et al., Blockade of VEGF-C signaling inhibits lymphatic malformations driven by oncogenic PIK3CA mutation. Nat Commun, 2020. 11(1): p. 2869.

41. Venot, Q., et al., Targeted therapy in patients with PIK3CA-related overgrowth syndrome. Nature, 2018. 558(7711): p. 540–546.

42. Adams, D.M., et al., Efficacy and Safety of Sirolimus in the Treatment of Complicated Vascular Anomalies. Pediatrics, 2016. 137(2): p. e20153257.

43. Delestre, F., et al., Alpelisib administration reduced lymphatic malformations in a mouse model and in patients. Sci Transl Med, 2021. 13(614): p. eabg0809.

44. Shin, Y., et al., Microfluidic assay for simultaneous culture of multiple cell types on surfaces or within hydrogels. Nat Protoc, 2012. 7(7): p. 1247–59.

45. Borikova, A.L., et al., Rho kinase inhibition rescues the endothelial cell cerebral cavernous malformation phenotype. J Biol Chem, 2010. 285(16): p. 11760–4.

46. Stockton, R.A., et al., Cerebral cavernous malformations proteins inhibit Rho kinase to stabilize vascular integrity. J Exp Med, 2010. 207(4): p. 881–96.

47. McDonald, D.A., et al., Fasudil decreases lesion burden in a murine model of cerebral cavernous malformation disease. Stroke, 2012. 43(2): p. 571–4.

48. Lekwuttikarn, R., et al., Genotype-Guided Medical Treatment of an Arteriovenous Malformation in a Child. JAMA Dermatol, 2019. 155(2): p. 256–257.

49. Edwards, E.A., et al., Monitoring Arteriovenous Malformation Response to Genotype-Targeted Therapy. Pediatrics, 2020. 146(3).

50. Homayun-Sepehr, N., et al., KRAS-driven model of Gorham-Stout disease effectively treated with trametinib. JCI Insight, 2021. 6(15).

51. Jacinto, E., et al., Mammalian TOR complex 2 controls the actin cytoskeleton and is rapamycin insensitive. Nat Cell Biol, 2004. 6(11): p. 1122–8.

52. Feldman, M.E., et al., Active-site inhibitors of mTOR target rapamycin-resistant outputs of mTORC1 and mTORC2. PLoS Biol, 2009. 7(2): p. e38.

53. Janes, M.R., et al., Effective and selective targeting of leukemia cells using a TORC1/2 kinase inhibitor. Nat Med, 2010. 16(2): p. 205–13.

54. Farhan, M.A., et al., Endothelial Cell mTOR Complex-2 Regulates Sprouting Angiogenesis. PLoS One, 2015. 10(8): p. e0135245.

55. Couto, J.A., et al., A somatic MAP3K3 mutation is associated with verrucous venous malformation. Am J Hum Genet, 2015. 96(3): p. 480–6.

56. Couto, J.A., et al., Somatic MAP2K1 Mutations Are Associated with Extracranial Arteriovenous Malformation. Am J Hum Genet, 2017. 100(3): p. 546–554.

57. Al-Olabi, L., et al., Mosaic RAS/MAPK variants cause sporadic vascular malformations which respond to targeted therapy. J Clin Invest, 2018. 128(4): p. 1496–1508.

58. Nikolaev, S.I., et al., Somatic Activating KRAS Mutations in Arteriovenous Malformations of the Brain. N Engl J Med, 2018. 378(3): p. 250–261.

59. Machesky, L.M. and A. Hall, Role of actin polymerization and adhesion to extracellular matrix in Rac- and Rho-induced cytoskeletal reorganization. J Cell Biol, 1997. 138(4): p. 913–26.

60. Hall, A., Rho GTPases and the actin cytoskeleton. Science, 1998. 279(5350): p. 509–14.

61. Liu, G.Y. and D.M. Sabatini, mTOR at the nexus of nutrition, growth, ageing and disease. Nat Rev Mol Cell Biol, 2020. 21(4): p. 183–203.

62. Li, W., et al., Hypoxia-induced endothelial proliferation requires both mTORC1 and mTORC2. Circ Res, 2007. 100(1): p. 79–87.

63. Wang, S., et al., Regulation of endothelial cell proliferation and vascular assembly through distinct mTORC2 signaling pathways. Mol Cell Biol, 2015. 35(7): p. 1299–313.

64. Tsuji-Tamura, K. and M. Ogawa, Inhibition of the PI3K-Akt and mTORC1 signaling pathways promotes the elongation of vascular endothelial cells. J Cell Sci, 2016. 129(6): p. 1165–78.

65. Huang, W., et al., mTORC2 controls actin polymerization required for consolidation of long-term memory. Nat Neurosci, 2013. 16(4): p. 441–8.

66. Thomanetz, V., et al., Ablation of the mTORC2 component rictor in brain or Purkinje cells affects size and neuron morphology. J Cell Biol, 2013. 201(2): p. 293–308.

67. Mendoza, M.C., E.E. Er, and J. Blenis, The Ras-ERK and PI3K-mTOR pathways: cross-talk and compensation. Trends Biochem Sci, 2011. 36(6): p. 320–8.

68. Ren, A.A., et al., PIK3CA and CCM mutations fuel cavernomas through a cancer-like mechanism. Nature, 2021. 594(7862): p. 271–276.

69. Snellings, D.A., et al., Developmental venous anomalies are a genetic primer for cerebral cavernous malformations. Nat Cardiovasc Res, 2022. 1: p. 246–252.

