## Supplementary material for "Microphysiological vascular malformation model reveals a role of dysregulated Rac1 and mTORC1/2 in lesion formation": Methods

### 2. METHODS

**Cell culture.** Human umbilical vein endothelial cells (HUVECs, Lonza) were cultured in growth medium (EGM-2, Lonza). Human lung fibroblasts (HLF, Lonza) were cultured in growth medium (FGM-2, Lonza), and HEK-293T cells (Clonetech) were grown in high glucose DMEM (Hyclone) supplemented with 10% fetal bovine serum (Hyclone) and 1% penicillin/streptomycin (Life Technologies). All cells were cultured at 37 °C and 5% $CO_2$ in a humidified incubator.

**Antibodies and cell biology reagents.** Anti-VE-cadherin (F-8) was from Santa Cruz Biotechnology. DAPI, rhodamine phalloidin (1 mg/mL), AlexaFluor-647 phalloidin (1 mg/mL), and AlexaFluor conjugated secondary antibodies were from Life Technologies. For antibody concentrations, see Immunofluorescence and Western Blot sections below. Rapamycin, Alpelisib (BYL719), Y-27362 2HCl, Trametinib (GSK1120212), Batimastat (BB-94), EHT 1864 2HCl, and PF-3758309 were from Selleckchem.

**Cloning and lentiviral infection.** Plasmids encoding pHAGE-EF1aL-eGFP-W (Addgene plasmid #126686, a gift from Darrell Kotton), pHAGE-PIK3CA-E542K-ires-GFP, pHAGE-PIK3CA-E545K-ires-GFP (Addgene plasmid #116479 and #116485, gifts from Gordon Mills and Kenneth Scott), pHAGE-PIK3CA-WT-ires-GFP, and pLenti.PGK.LifeAct-Ruby.W (Addgene plasmid #51009, a gift from Rusty Lansford) were used for lentivirus generation. Q5 site directed mutagenesis (NEB) was performed on pHAGE-PIK3CA-E542K-ires-GFP to generate pHAGE-PIK3CA-WT-ires-GFP. For lentivirus production, HEK-293T cells were transfected with pHAGE-plasmid, pSPAX2 plasmid and pMD2.G plasmid (Addgene plasmid #12260 and #12259, gifts from Didier Trono) using calcium phosphate transfection. Lentiviral supernatants were collected 2 days after transfection and concentrated using PEG-IT (System Biosciences, Palo Alto, CA) viral precipitator. Concentrated lentivirus was resuspended in PBS. Serial dilutions of lentiviral particles were performed to determine the optimal concentration for transduction. Concentrated lentiviruses were subsequently added to 100,000 HUVECs cultured in 6 well plate. Transduction efficiency was determined by GFP immunofluorescence. Transduced HUVECs were used between passages 5-11.

**PDMS wells for network formation.** To generate wells for vascular networks, polydimethylsiloxane (PDMS, Sylgard 184, Dow-Corning) was mixed at 10:1 base:curing agent, poured into the lid of a 150 mm culture dish, and degassed in a dessicator under house vacuum for 1 hr. PDMS was cured overnight at 60 °C. Cured PDMS was then removed from the lid, and a 6-mm biopsy punch and razor were used to create 1 cm x 1 cm squares of PDMS with a 6-mm diameter hole at the center. Clear tape and isopropanol were used to clean the PDMS surfaces and glass coverslip. PDMS and coverslips were treated with oxygen plasma (Harrick Plasma) for 30s, bonded, and incubated at 100 °C for 20 min. 0.01% (w/v) poly-L-lysine in water (Sigma) was added to the interior of the wells and incubated at RT for 5 min before 3x washes with sterile, distilled (DI-) $H_2O$. 1% (w/v) glutaraldehyde in water was then added to wells and incubated at RT for 5 min before 3X washes with sterile DI-$H_2O$ and incubation overnight in DI-$H_2O$ on a laboratory orbital shaker overnight at RT. Clean, surface treated wells were then wrapped in aluminum foil, autoclaved, dried at 100 °C for 20 min, and placed in sterile wells of a 24-well plate.

**Hydrogel formation and seeding in PDMS wells.** Fresh solutions of fibrinogen from bovine plasma (Millipore Sigma) were prepared prior to each seeding by dissolving fibrinogen powder in Dulbecco's Phosphate Buffered Saline (DPBS, Millipore Sigma) to a concentration of 5 mg/mL and incubating at 37 °C for 30 min prior to filter sterilization with 0.2 µm syringe filter

(Millipore Sigma). Thrombin from bovine plasma was solubilized in DPBS to a concentration of 100 U/mL, aliquoted, frozen at -80 °C, and thawed immediately before each seeding. HUVECs and HLFs were lifted from culture flasks with 0.25% (w/v) Trypsin-EDTA (ThermoFisher), centrifuged at 200 x *g* for 5min. HUVECs and HLFs were each resuspended at $13x10^6$ cells/mL of EGM-2. For each well, 50µL of cells in fibrin pre-polymer were prepared by mixing 25µL of 5 mg/mL fibrinogen, 11.5µL of $13x10^6$ HUVECs/mL EGM-2, 11.5µL of $13x10^6$ HLFs/mL EGM-2, 0.5 µL of 100 U/mL thrombin, and 1.5µL of EGM-2 for final cell concentrations of $3x10^6$ cells/mL in the total slurry. After mixing with a pipette to avoid bubbles, a 50µL slurry was added to each PDMS well, incubated for 15 min at 37 °C in a cell culture incubator, with inversion every few minutes to allow cells to be distributed in 3D. 2.5mL of fresh EGM-2 was added into each well of the 24-well plate containing a PDMS well, and media was changed daily.

**Microfluidic device fabrication.** Microfluidic devices were fabricated using a protocol adapted from previous approaches [1-3]. Briefly, film masks (Fineline) were printed from technical drawings made with AutoCAD (Autodesk). A mask aligner (SUSS MicroTec) was used to expose negative photoresist (SU8-2150, Microchem) spun onto 4-in silicon wafers (University Wafers) at a thickness of 200 mm to 350 nm wavelength light (1500 mJ/$cm^2$). After post-exposure bakes at 60 °C and 100 °C, uncrosslinked photoresist stripped using propylene glycol monomethyl ether acetate (PGMEA, Millipore Sigma) developer and dried with isopropanol. After plasma treatment for 30 s, trichloro(1H,1H,2H,2H-perfluorooctyl)silane (Sigma) was vapor deposited onto the patterned silicon wafers (referred to subsequently as silicon masters). Individual devices were then fabricated using standard soft lithography. PDMS (10:1 base:curing agent) was poured onto silicon masters, degassed using house vacuum, and cured at 60 °C overnight. Cured PDMS was then removed from the master, cut and trimmed with a razor, and media and hydrogel ports were punched out with biopsy punches (6mm and 1.5mm, respectively). The surface of the PDMS were cleaned with tape and isopropanol. Cleaned PDMS devices and glass coverslips were then treated with oxygen plasma for 30 s, bonded, and incubated at 100 °C for 20 min. Prior to use, devices were sterilized with UV treatment for 20 min.

**Hydrogel formation and seeding in microfluidic devices.** Hydrogels were prepared following the same protocol as the PDMS wells with the exception of different cell concentrations for each channel. For the HUVEC channel, HUVECs were resuspended at a concentration of $5.5x10^6$ HUVECs/mL in EGM-2 media and added to an equal volume of 5 mg/mL fibrinogen diluted in PBS. To visualize ECM degradation and vascular lesion formation, 1.5mg/mL of AlexaFluor-64-conjugated fibrinogen (ThermoFisher) was added to the hydrogel. After mixing, 1U of thrombin was added to the slurry and mixed prior to injection into the device. Devices were then flipped upside down and incubated for 15 minutes at 37 °C in a cell culture incubator, inverting every few minutes to ensure even distribution of cells in 3D. The same protocol was used for preparing HLF-fibrin hydrogels for the HLF channels with the exception that the HLF suspension was prepared at a density of $2.75x10^6$ HLFs/mL in EGM-2 media. Media was then changed daily and devices were fixed 3 days after seeding. For mitotic inhibition, HUVECs plated onto 10cm dishes were incubated with 0.01mg/mL mitomycin-C (source) for 2.5 hours at 37 °C. Mitomycin-C treated HUVECs were trypisinized and seeded into microfluidic devices as described above.

**Immunofluorescence.** To observe network assembly over time, PDMS wells loaded with HUVECs expressing GFP (see Cloning and lentiviral infection, above) were imaged daily with a laser scanning confocal microscope (FV3000, Olympus) with a 488 nm laser diode. Z-stacks of the gel volume were taken at 10x magnification (U Plan S-Apo, 0.4 NA air objective, Olympus). For endpoint immunostaining, PDMS wells or devices were fixed with 4% paraformaldehyde (Millipore Sigma) in PBS containing calcium and magnesium (PBS++) at 37 °C for 15 min. After rinsing twice with PBS++, devices were left on a laboratory rocker for 24 hrs in PBS++ to wash. Cells were then permeabilized with 0.3% Triton X-100 (Millipore Sigma) for 10 min at RT and nonspecific antibody binding was blocked with 2% (w/vol) BSA in PBS++ for 24 hrs at RT. Wells and devices were then sealed in tissue culture plates and kept at 4 °C until antibodies were added. Samples cultured in microfluidic devices were imaged daily at 4x magnification (U Plan Fluor, 0.13 NA air objective, Olympus) on a widefield microscope and fixed 3 days after seeding. For immunostaining, cells were permeabilized with 0.3% Triton X-100. Primary antibodies against VE-cadherin (mouse, anti-human F-8; Santa Cruz Biotechnology) were diluted in 2% BSA in PBS++ (1:200, vol/vol) and applied overnight on a laboratory rocker at 4 °C. Cells were then rinsed three times over 1 hr with PBS++, and secondary antibodies (goat, anti-mouse IgG conjugated to AlexaFluor-647; Thermo-Fisher Scientific) were diluted in 2 % BSA in PBS++ (1:200, vol/vol) and applied to devices at room temperature for 1 hr on a laboratory rocker before rinsing three times over 30 minutes with PBS++. F-actin was labeled with AlexaFluor-488 or rhodamine-phalloidin (Thermo Fisher), and the nucleus was labeled with DAPI (Thermo Fisher) diluted in PBS++ (1:200 and 1:1000, vol/vol, respectively) for 1 hr at room temperature before rinsing three times over 30 min with PBS++. Images were acquired with an Olympus FV3000 laser scanning confocal with a 30x U Plan S-Apo N 1.05 NA silicone oil immersion objective, and images were adjusted for brightness and contrast using ImageJ. For computational image processing used to measure cell and network topology, see Supplementary Material.

**Western blot and small GTPase activity assays**
Cells were cultured in complete media unless otherwise noted. For measuring changes in phospho-AKT and phospho-ERK level in response to Alpelisib, Rapamycin, or Trametinib, HUVECs were cultured in reduced serum growth medium (0.5% FBS) overnight and then changed into media containing 2% FBS with or without inhibitor. After 10 minutes, cells were lysed on ice with RIPA buffer (ThermoFisher Scientific) containing Halt protease and phosphatase inhibitor (ThermoFisher Scientific). Clarified lysates were resolved on Novex 4-12% Bis-Tris gel (ThermoFisher Scientific) and transferred to a PVDF membrane. Standard immunoblotting protocols were performed for western blotting with primary antibodies used at the following concentrations: anti-pAKT-Ser473 (1:1000, Cell signaling #9271), anti-AKT (1:1000, Cell Signaling #9272), anti-pERK1/2-Thr202/Tyr204 (1:1000, Cell Signaling #9101), anti-ERK1/2 (1:1000, Cell Signaling), and anti-GAPDH (1:2000, Cell Signaling #2118). HRP-conjugated secondary antibodies and SuperSignal West Femto chemiluminescent substrate were used for detection. Western blot images were quantified with FIJI/ImageJ. Proteome profiler human protease array (RnDSystems) was performed according to manufacturer instructions.

Ras, Rac1, and Cdc42 pull-down assays (Cytoskeleton Inc) were performed according to the provided instruction manual. Briefly, confluent cells cultured overnight in reduced serum conditions were stimulated with complete media with or without inhibitor for 10 minutes. Cells were rinsed once with cold PBS++ and lysed with cold lysis buffer (Cytoskeleton Inc, 50mM Tris pH 7.5, 10mM MgCl2, 0.3M NaCl, 2% IGEPAL, and protease and phosphatase inhibitor cocktails). Lysates were sonicated at 3W on ice for 10s, and clarified at 14,000g for 5 min. Protein concentration was quantified with BCA assay (ThermoFisher Scientific), and protein concentration and volume were equalized with cell lysis buffer. 500 µg of lysate was incubated with 10 µg of PAK-PBD or RAF-RBD beads on a rotator for an hour at 4 °C. Bead pellets were washed three times with wash buffer (Cytoskeleton Inc, 25mM Tris pH 7.5, 30mM MgCl2, 40mM NaCl) and extracted with 2x NuPAGE LDS containing 100mM DTT. For quantification, active RAC1 and RAS1 were normalized to total RAC1 and RAS1, respectively.

**Live cell imaging, migration tracking and quantification**
HUVECs transduced with LifeAct-mRuby were cultured overnight on 35mm dishes (Ibidi). Live imaging of microvascular networks was performed with an Olympus FV3000 resonant scanning confocal system equipped with a stage-top incubator. Images were taken at 10x magnification (U Plan S-Apo, 0.4 NA air objective, Olympus), every 5 minutes for a period of 72 hours. Live imaging of cell migration was carried out on the same system but with a Plan S-Apo 30x/1.05 NA Silicone Oil (Olympus) objective. Images were captured every minute for two hours. Tracking of endothelial cells was performed using the cellpose TrackMate[4-6] module on Fiji. A pretrained cytoplasm model with an estimated 30μm cell diameter was used for segmenting cells. To confirm tracking accuracy, cell migration trajectories were visually inspected. Migration tracks were plotted with Matlab. Mean migration speed and confinement ratios were plotted and statistical analyses were performed using GraphPad Prism. Mean square displacement plot was generated using MotilityLab[7].

**Beta-galactosidase staining**
HUVECs were plated onto 12 well plate at a density of 25,000 cells per well. Cells were fixed 24 hours after seeding and stained for beta-galactosidase activity (Cell Signaling #9860S) according to manufacturer's protocol. Images were captured with widefield microscopy. Cells with positive beta-gal staining were quantified using manual thresholding in FIJI/ImageJ. The same threshold value was used for all images.

**Junctional and actin quantification.**
Adherens junction and actin content were quantified using methods previously described, with modifications [8]. To quantify adherens junction structure and assembly of HUVECs cultured in 2D, greyscale images of HUVECs immunostained with VE-cadherin were binarized on a threshold based on the mean cytosolic intensity of VE-cadherin for each image. A region of interest (ROI) was drawn within the cytosol of a cell for each image, the mean intensity within the ROI was determined, and the threshold for binarization was set to 2X this mean intensity value. Hot pixel noise was removed using the "Despeckle" command in ImageJ. To remove perinuclear signal, the DAPI channel for each image was binarized, dilated 3X, and inverted to create a mask that was then applied to the despeckled, binarized VE-cadherin images. Junctional area was defined as the total number of nonzero pixels. To quantify cortical actin, the intensity of Alexa Fluor 488 phalloidin-labelled cells was plotted along lines drawn between the centroids in the nuclei of neighboring cells. Local intensity peaks were measured to determine the location of stress fibers and cortical actin. Cortical actin was defined as the area under the peak at cell–cell junctions normalized to total area under the curve.

**Cell count and area quantification.**
Cell area was quantified using a custom Cell Profiler pipeline. Briefly, DAPI channel was used for segmenting individual cell nuclei for both cell count and area quantification. The binary mask containing the segmented nuclei was used as a reference for segmenting cells. Thresholding for cells was based on actin staining and was determined using a minimum cross entropy thresholding method.

**Quantification of vascular network topology and fibrin void regions.**
Confocal z-stacks (4 μm step size, captured from top to bottom of the HUVEC channel within microfluidic devices) were used for measurement of vascular network topology. Rolling z projections (average over five z-sections, FIJI) were performed to increase the signal to noise ratio in the images. Maximal intensity rolling z projections were performed for DAPI and Actin channels, and minimum intensity rolling z projection was performed for Fibrin ECM channel. Contrast enhancement (0.35% of saturated pixels) was used to normalize image brightness

across the z-stack. The processed z-stacks were exported as image sequences for 2D segmentation in Cell Profiler. DAPI and Actin channels were used for segmenting vascular networks. An adaptive minimum cross-entropy thresholding method (size of adaptive window = 250 pixels) was used for segmenting vascular networks. Briefly, a binary mask containing segmented nuclei was used as a reference for segmenting vascular networks labeled with actin. Binarized images were reconstructed into a z-stack and used as input images for 3D skeletonization, pruning, and vasculature analysis in VesselVio [9]. To quantify fibrin void regions, processed fibrin images were binarized using adaptive thresholding (size of adaptive window = 100 pixels, Sauvola thresholding method). Binarized images were inverted and reassembled into z-stacks. Volume filling and 3D void volume analysis of binarized fibrin void regions was performed in VesselVio[9].

**Statistical analysis.**
Graphs and statistical analyses were generated in Prism 9 (GraphPad). Data were plotted as individual points with mean ± sem. Unless otherwise mentioned, statistical differences between two sets of normally distributed data were calculated using unpaired, two-tailed Student's test. Statistical differences on more than two groups of data were determined using one-way ANOVA followed by Tukey posthoc test.

### References:


[1] S. Kim, H. Lee, M. Chung, N.L. Jeon, Engineering of functional, perfusable 3D microvascular networks on a chip, Lab on a Chip 13(8) (2013) 1489-1500.
[2] W.J. Polacheck, J.L. Charest, R.D. Kamm, Interstitial flow influences direction of tumor cell migration through competing mechanisms, Proc Natl Acad Sci U S A 108(27) (2011) 11115-20.
[3] W.J. Polacheck, M.L. Kutys, J.B. Tefft, C.S. Chen, Microfabricated blood vessels for modeling the vascular transport barrier, Nat Protoc 14(5) (2019) 1425-1454.
[4] J.Y. Tinevez, N. Perry, J. Schindelin, G.M. Hoopes, G.D. Reynolds, E. Laplantine, S.Y. Bednarek, S.L. Shorte, K.W. Eliceiri, TrackMate: An open and extensible platform for single-particle tracking, Methods 115 (2017) 80-90.
[5] D. Ershov, M.S. Phan, J.W. Pylvanainen, S.U. Rigaud, L. Le Blanc, A. Charles-Orszag, J.R.W. Conway, R.F. Laine, N.H. Roy, D. Bonazzi, G. Dumenil, G. Jacquemet, J.Y. Tinevez, TrackMate 7: integrating state-of-the-art segmentation algorithms into tracking pipelines, Nat Methods 19(7) (2022) 829-832.
[6] C. Stringer, T. Wang, M. Michaelos, M. Pachitariu, Cellpose: a generalist algorithm for cellular segmentation, Nat Methods 18(1) (2021) 100-106.
[7] I.M. Wortel, A.Y. Liu, K. Dannenberg, J.C. Berry, M.J. Miller, J. Textor, CelltrackR: an R package for fast and flexible analysis of immune cell migration data, ImmunoInformatics 1 (2021) 100003.
[8] W.J. Polacheck, M.L. Kutys, J. Yang, J. Eyckmans, Y. Wu, H. Vasavada, K.K. Hirschi, C.S. Chen, A non-canonical Notch complex regulates adherens junctions and vascular barrier function, Nature 552(7684) (2017) 258-262.
[9] J.R. Bumgarner, R.J. Nelson, Open-source analysis and visualization of segmented vasculature datasets with VesselVio, Cell Rep Methods 2(4) (2022) 100189.
